# In-depth phylogenomic analysis of arbuscular mycorrhizal fungi based on a comprehensive set of de novo genome assemblies

**DOI:** 10.1101/2021.04.09.439159

**Authors:** Merce Montoliu-Nerin, Marisol Sánchez-García, Claudia Bergin, Verena Esther Kutschera, Hanna Johannesson, James D. Bever, Anna Rosling

## Abstract

- Morphological characters and nuclear ribosomal DNA (rDNA) phylogenies have so far been the basis of the current classifications of arbuscular mycorrhizal (AM) fungi. Improved understanding of the phylogeny and evolutionary history of AM fungi requires extensive ortholog sampling and analyses of genome and transcriptome data from a wide range of taxa.
- To circumvent the need for axenic culturing of AM fungi we gathered and combined genomic data from single nuclei to generate *de novo* genome assemblies covering seven families of AM fungi. Comparative analysis of the previously published *Rhizophagus irregularis* DAOM197198 assembly confirm that our novel workflow generates high-quality genome assemblies suitable for phylogenomic analysis. Predicted genes of our assemblies, together with published protein sequences of AM fungi and their sister clades, were used for phylogenomic analyses.
- Based on analyses of sets of orthologous genes, we highlight three alternative topologies among families of AM fungi. In the main topology, Glomerales is polyphyletic and Claroideoglomeraceae, is the basal sister group to Glomeraceae and Diversisporales.
- Our results support family level classification from previous phylogenetic studies. New evolutionary relationships among families where highlighted with phylogenomic analysis using the hitherto most extensive taxon sampling for AM fungi.

## Introduction

Arbuscular mycorrhizal (AM) fungi are an ecologically important group of fungi that form ubiquitous associations with plants, establishing symbiosis with up to 80% of land plants species (Parniske, 2008; Smith and Read, 2008). AM fungi play foundational roles in terrestrial productivity, and there is accumulating evidence that AM fungal taxa are functionally distinct and that their community composition have functional consequences for terrestrial ecosystems (Hoeksema *et al*., 2018; Koziol *et al*., 2018). Therefore, progress in understanding the ecologically distinct roles of AM fungi depends upon accurate phylogenetic inference at all taxonomic levels.

Available literature identify that all AM fungi form a monophyletic lineage within the fungal kingdom. This lineage is taxonomically classified either as a phylum, Glomeromycota, (Schüβler, Schwarzott and Walker, 2001; James *et al*., 2006; Hibbett *et al*., 2007; Schüßler and Walker, 2010; Tedersoo *et al*., 2018) or as the sub-phylum Glomeromycotina, which together with Morteriellomycotina and Mucoromycotina, make up the phylum Mucoromycota (Spatafora *et al*., 2016; James *et al*., 2020, Li *et al*., 2021). The current consensus classification of AM-fungal species into genera and families was established by Redecker and co-authors in 2013, when systematists with long experience in the biology and taxonomy of AM fungi joined forces to integrate morphological and molecular phylogenetic evidence to generate a meaningful classification that reflects evolutionary relationships (Redecker *et al*., 2013). Molecular evidence at the time was primarily based on partial sequences of nuclear ribosomal DNA (rDNA). Around 300 species of AM fungi are currently described and classified into 33 genera, twelve families and four orders (Redecker et al., 2013; Wijayawardene *et al*., 2018).

Molecular identification of AM fungi to genera and family is usually possible based on sequences of small subunit (SSU) or large subunit (LSU) regions of rDNA genes (Redecker *et al*., 2013, Öpik *et al*., 2013). However, species level inference based on rDNA genes is difficult due to high levels of intra species variation (Stockinger, Walker and Schüßler, 2009; House *et al*., 2016). While the rDNA operon is commonly found in a multi-copy tandem repeat organization across fungi (Lofgren *et al*., 2019), in AM fungi different rDNA variants can be scattered across the genome (Vankuren *et al*., 2013; Maeda *et al*., 2018) and lack the usual tandem organization (Maeda *et al*., 2018). The fact that rDNA genes are present as paralogs in AM fungal genomes (Maeda *et al*., 2018) likely explains the high levels of within strain and species diversity of rDNA sequences.

Limitations of single-locus phylogenetic inference and paralogous nature of rDNA genes in AM fungi, calls for the need to generate extensive ortholog datasets from taxa representing different families, in order to accurately infer phylogenetic relationships among AM fungal lineages. One approach in this direction was achieved in a recent study using spore transcriptomic data for phylogenomic analysis of nine taxa from seven families (Beaudet *et al*., 2018). In the study Glomerales was recovered as polyphyletic (Beaudet *et al*., 2018), in contrast to earlier ribosomal genes phylogenies where Glomerales was found to be monophyletic (Krüger *et al*., 2012). Other phylogenomic studies have not adressed relations among families largelly due to limited taxon sampling (Sun *et al*., 2018; Morin *et al*., 2019, Venice *et al*., 2020, Li *et al*., 2021). Due to difficulties in obtaining enough pure DNA for whole genome sequencing available genomic data still represent only a fraction of the diversity of AM fungi.

Progress in AM fungal genomics has been limited by their biology. AM fungi complete their life cycle underground, as obligate symbionts of plant roots, and reproduce through multinuclear asexual spores (Bonfante and Genre, 2010). The spores are the largest isolable structure produced, but large-scale isolation of spores is needed in order to obtain enough DNA for whole genome sequencing. Such large-scale harvest of spores is only possible with AM fungi grown in axenic cultures where the fungus produces spores in a compartment separate from the transformed plant roots that it associates with (Tisserant *et al*., 2013), or for the rare taxa, such as *Diversispora epigaea* that forms fruitbodies above ground, from which large amounts of spores can be extracted (Sun *et al*., 2018). Axenic culturing methods are time-consuming and have only been successful for a handful of species (Kameoka *et al*., 2019). Due to the difficulty of producing clean cultures and isolate high quality DNA extracts for the majority of AM fungal species, it has been a slow path towards genomic studies of AM fungi (Tisserant *et al*., 2013; Lin *et al*., 2014; Chen *et al*., 2018; Kobayashi *et al*., 2018; Sun *et al*., 2018; Morin *et al*., 2019, Venice *et al*., 2020).

AM fungal hyphae and their asexual spores are coenocytic, and sequence analysis of individual nuclei have been used to analyze intra organismal polymorphism mainly by mapping reads of single nuclei to reference genomes of the model AM fungus *Rhizophagus irregularis* (Lin *et al*., 2014; Ropars *et al*., 2016; Chen *et al*., 2018). To circumvent the obstacle of pure culturing, a workflow was developed that takes advantage of automated nuclei sorting by extracting nuclei from un-germinated spores, directly extracted from soil. Single nuclei were sorted, followed by whole genome amplification (WGA) and sequencing. Finally, the data from several nuclei were combined to build a *de novo* genome assembly of *Claroideoglomus claroideum* (Montoliu-Nerin *et al*., 2020). With this novel workflow AM fungal genome assemblies can be obtained from as little as one single spore, independently of the species ability to grow in axenic cultures. Similar approaches have been successfully applied in other organisms for which limited access to pure biological material suitable for extraction of high-quality DNA has prevented genome sequencing (Stepanauskas and Sieracki, 2007; Woyke *et al*., 2009; Heywood *et al*., 2011; Yoon *et al*., 2011; Walker *et al*., 2014; Wideman *et al*., 2019).

In this study, we sorted and sequenced nuclei from AM fungal spores representing species across Glomeromycota, aiming to have two taxa for each genus. To evaluated the quality of assemblies generated by our workflow, we included *Rh. irregularis* DAOM197198, a strain for which a reference genome generated from an axenic culture is available (Chen *et al*., 2018), and compared this published assembly with our newly generated assembly. A final count of 22 strains, from ten genera, across seven families were successfully sequenced, and *de novo* genome assemblies were constructed. Our newly generated dataset contains 18 taxa for which genome data was previously not available. This comprehensive taxon sampling allowed us to infer evolutionary relationships among AM fungi. Furthermore, the release of new whole genome assemblies provides a resource to the research community, for those interested in further exploring genetics and evolution of this important group of fungi.

## Material and methods

### Fungal strains

Taxa were selected to represent most families in Glomeromycota (Schüßler, & Walker, 2010; Redecker et al., 2013), aiming for two species per genus (Table S1). The isolates were obtained as whole inoculum from the International culture collection of (vesicular) arbuscular mycorrhizal fungi (INVAM) at West Virginia University, Morgantown, WV, USA (*Acaulospora colombiana, Acaulospora morrowiae, Ambispora gerdemannii, Ambispora leptoticha, Cetraspora pellucida, Claroideoglomus candidum, Claroideoglomus claroideum, Dentiscutata erythropa, Dentiscutata heterogama, Diversispora eburnea, Funneliformis caledonius, Gigaspora rosea, Racocetra fulgida, Racocetra persica, Scutellospora calospora, Paraglomus brasilianum, Paraglomus occultum*), James D. Bever’s lab, University of Kansas, USA (*Cetraspora pellucida, Claroideoglomus candidum, Funneliformis mosseae, Gigaspora margarita*), and as spores in a tube from Agriculture and Agri-food Canada, Government of Canada (*Rh. irregularis* DAOM197198). In addition to the AM fungi sampled for this study, we included the annotated genome assemblies of previously sequenced isolates of AM fungi (Table S2). Furthermore, we included all species of the closest sister lineages with available genome assemblies and annotations (December 2019) in the JGI (Joint Genome Institute) database, i.e. Morteriellomycota (2 taxa, Mondo *et al*., 2017; Uehling *et al*., 2017) and Mucoromycota (12 taxa, Ma *et al*., 2009; Wang *et al*., 2013; Schwartze *et al*., 2014; Chibucos *et al*., 2016; Corrochano *et al*., 2016; Mondo, Dannebaum, *et al*., 2017; Mondo, Lastovetsky, *et al*., 2017; Chang *et al*., 2019) (Table S2). Finally, representatives of Dikarya was included as an outgroup, with three representatives of Ascomycota, one taxon each from the subphyla Taphrinomycotina (Pomraning *et al*., 2018), Saccharomycotina (Wood *et al*., 2002), and Pezizomycotina (Martin *et al*., 2010); and three representatives of Basidiomycota, one taxon each from the subphyla Agaricomycotina (Martin *et al*., 2008), Ustilaginomycotina (Kämper *et al*., 2006), and Pucciniomycotina (Schwessinger *et al*., 2018) (Table S2).

### Nuclear sorting and whole genome amplification

Spores were extracted from whole inoculum cultures by sieving, followed by a sucrose gradient centrifugation as described in Montoliu-Nerin *et al*., 2020. A single spore or a pool of spores (Table S1) were then rinsed and stored in 20 µl ddH2O in a 1.5 ml tube. Spores were crushed with a sterile pestle after adding 180 µl of 1x PBS and DNA was stained by adding 1 µl of 200x SYBR Green I Nucleic Acid stain (Invitrogen™, Thermo Fisher Scientific, MA, USA). The nuclear sorting with flow activated cell sorter (FACS) proved to be more successful when the crushed spore solution was transferred to the small 0.5 ml tube for staining. This allowed the spore debris to settle while the nuclei remained in solution. The sample was left staining for 30 to 60 minutes, and lower sorting performance was observed when exceeding that time. The nuclear sorting was performed at the SciLifeLab Microbial Single Cell Genomics Facility with a MoFlo™ Astrios EQ sorter (Beckman Coulter, USA), as in Montoliu-Nerin *et al*., 2020. Briefly, a 100 um nozzle was used and the sheath fluid, 0.1 um filtered 1x PBS, was run at 25 psi. Nuclei populations were identified via nuclei acid staining using the 488 nm laser and a 530/40 bandpass filter over forward or side scatter. Individual nuclei were deposited into 96- or 384-well plates using stringent single-cell sort settings (single mode, drop envelope 1). These sort-settings abort target cells if another particle of any type is in the same or the neighboring drop, thereby increasing the number of aborts while ensuring that only one particle gets sorted per well. Each day of sorting, the sort precision was determined with beads sorted onto a slide and counted manually under the microscope. A low event rate was used to decrease the risk of sorting doublets, for most samples below 500 events per second with a drop generation of >40,000 per second corresponding to well below 1% of nuclei in the samples. Most of the remaining particles were low in SYBR Green fluorescence. To each plate, up to 48 nuclei were sorted leaving the rest of the wells empty. Plates with sorted nuclei were stored at -80°C.

DNA from the nuclei was amplified with the enzyme Phi29 with multiple displacement amplification (MDA). MDA reactions were run under clean conditions using the RepliPhi kit (Epicentre) in a 15 ul reaction volume in 96-well plates or with the Repli-g Single Cell kit (Qiagen) in a 10 ul reaction volume in 384-well plates. The nucleic acid stain SYTO 13 was added to the reaction to follow the DNA amplification over time. Protocol including plate size and MDA kit was changed over time (Table S1, detailed soring information is available in the linked public OSF and Supplementary data file 1).

### Sequencing of single nuclei

Amplified DNA of single nuclei were screened for the presence of DNA of fungal or bacterial origin through PCR amplification of rDNA markers using fungal and bacterial specific primers, following the protocol in Montoliu-Nerin *et al*., 2020. Depending on the success rate of sorting and MDA, a total of 7 to 24 nuclei from each isolate were sequenced independently with Illumina HiSeq-X, at the SNP&SEQ Technology Platform in Uppsala at the National Genomics Infrastructure (NGI) Sweden and Science for Life Laboratory, as in Montoliu-Nerin *et al*., 2020, changing to the TruSeq PCRfree DNA library preparation kit (Illumina Inc.) when enough DNA was available.

### Genome assembly and strain verification

Whole genome assembly was performed according to assembly workflow 3 as described in Montoliu-Nerin *et al*., 2020, in which all sets of reads from individually sequenced nuclei from each strain were combined and normalized using bbnorm of BBMap v.38.08 (Bushnell, 2014), setting an average depth of 100X, and then assembled using SPAdes v.3.12.0 (Bankevich *et al*., 2012). We chose workflow 3 for this study as it gives a good representation and accuracy of single copy genes, making it more suitable for downstream phylogenomic analyses than the other two workflows tested (Montoliu-Nerin *et al*., 2020). We used Quast v.4.5.4 (Gurevich et al., 2013) to quantitively assess the assemblies (Table S3) and ran BUSCO v.3.0.2b (Simão et al., 2015) to assess completeness of the genome, using fungi_odb9 as lineage setting, and rhizopus_oryzae as species set (Table S3-4). Raw reads and *de novo* genome assemblies are deposited in ENA under the accessions xx-yy.

To verify strain identity based on a reconstructed ribosomal gene phylogeny, we extracted the ribosomal gene operon from each newly assembled genome. For strains in the family Claroideoglomeraceae only one of its highly diverging rDNA sequences (Vankuren *et al*., 2013) were retrieved as a complete operon. We have previously identified both rDNA variants in *Claroideoglomus claroideum* using a different assembly workflow (Montoliu-Nerin *et al*., 2020). As demonstrated in that study, assembly workflow 3, which was used to produce an assembly with a better representation of single copy orthologs, failed to assemble the second rDNA variant. The SSU region was combined with the SSU alignment from Krüger *et al*., (2012). The whole rDNA operons extracted from the genome assemblies with verified identity, were aligned and a phylogeny was reconstructed with RAxML v.8.2.10 (Stamatakis, 2014), implementing the GTR model and with IQ-TREE v1.6.5 (Nguyen et al. 2015), using ModelFinder (Kalyaanamoorthy et al., 2017) and searching for the best partitioning scheme. Both analyses were ran with 1000 bootstrap replicates. Extracted rDNA operons for all *de novo* genome assemblies are deposited in ENA under the accessions xx-yy.

### Genome annotation

Each genome assembly was annotated using a snakemake workflow (Köster and Rahmann, 2012) v.2.0. The workflow is publicly available at https://bitbucket.org/scilifelab-lts/genemark_fungal_annotation/ (tag v3.0), with minor updates providing the same functionality. Briefly, repeats and transposable elements were *de novo* predicted in each of the assemblies using RepeatModeler v1.0.8 (Smit and Hubley, 2015) and the resulting repeat library was used to mask each genome assembly using RepeatMasker v4.0.7 (Smit *et al*., 2015).

UniProt/Swiss-Prot (Bateman *et al*., 2017) protein sequences (downloaded 8 May 2018) were aligned to each of the repeat-masked genome assemblies with MAKER v3.01.1-beta (Cantarel *et al*., 2008). Protein coding genes were *de novo* predicted from each of the repeat-masked genome assemblies with GeneMark-ES v4.33 (Ter-Hovhannisyan *et al*., 2008), providing the genomic locations of Uniprot/Swiss-Prot proteins aligned to the genome assembly to guide the gene predictions. Minimum contig size to be included in self-training of the GeneMark gene prediction algorithm was calculated to include at least 10Mb of training data, depending on the level of fragmentation of the assembly, and was set accordingly using the parameter “--min_contig” (Table of specific parameter used for each assembly is available in the linked public OSF). Protein and gene names were assigned to the gene predictions using a BLASTp v2.7.1 (Camacho *et al*., 2009) search of predicted protein sequences against the UniProt/Swiss-Prot database with default e-value parameters (1×10^−5^). InterProScan v5.30-69.0 (Cock *et al*., 2013) was used to collect predictive information about the predicted proteins’ functions.

### Assessing assembly quality

To confirm the accuracy of our workflow we included the reference strain *Rh. irregularis* DAOM197198 (Table S1) and compared the generated *de novo* genome assembly to a published high-quality genome assembly DAOM197198 v.2.0 (Table S2) (Chen *et al*., 2018). Including this well characterized strain allowed us to assess the performance of our assembly workflow described above. To assess efficiency and coverage of single nuclei MDA and sequencing, we mapped reads from individual nuclei back to the published reference assembly and to our *de novo* assembly of *Rh. irregularis* DAOM197198, using BWA 0.7.15 (Li and Durbin, 2009), and measured both % of reads mapping and % of assembly covered with mapped reads using Qualimap 2.2.1 (Okonechnikov, Conesa and García-Alcalde, 2016) and bamtools 2.3.0 stats (Barnett *et al*., 2011). We also tested for polymorphism introduced during MDA by pair-wise alignment of the 271 BUSCO genes retrieved from both assemblies of *Rh. irregularis* DAOM197198 using MAFFT 7.407 (Katoh and Standley, 2013) Percent similarity for the alignments was calculated with esl-alistat in HMMer 3.2.1 (Hancock *et al*., 2004). Finally, orthogroups were identified with OrthoFinder 2.4.0 (Emms and Kelly, 2018) using standard settings.

### Phylogenomic analyses

Phylogenomic analyses were performed at different taxonomic scales, using datasets with different taxon sampling (Table S2-S3). For each set of taxa, single copy orthologs (SCOs) were identified from the gene predictions using OrthoMCL v.2.0.9 (Li *et al*., 2003) using default parameters. SCOs present in >50% or 100% of the taxa were selected. Amino acid sequences were aligned using MAFFT v.7.407 (Katoh and Standley, 2013). Poorly aligned regions were removed using trimAl v.1.4.1 (Capella-Gutiérrez *et al*., 2009) with a gap threshold of 0.1 (0.2 in the dataset with only 15 taxa selected, Table S4). Individual SCO alignments were removed if shorter than 100 amino acids. SCO alignments were used either separately, to produce individual gene trees, or concatenated, to produce maximum likelihood (ML) consensus trees. Individual SCO alignments were concatenated into a supermatrix using the script geneStitcher.py (Schluter, 2016), which also produces a partition file, with one partition per gene. Lists of SCOs used for phylogenomic inferences for each set of taxa and their corresponding gene annotations are available in the public OSF repository.

Phylogenomic inferences based on the concatenated sets of SCOs were performed using two maximum likelihood (ML) methods. All ML phylogenies were inferred using RAxML v.8.2.10 (Stamatakis, 2014), with 100 bootstrap replicates, and with a partitioned model that treated each SCO as a separate partition and implementing the PROTOGAMMAWAG model for all partitions. Secondly, ModelFinder (Kalyaanamoorthy *et al*., 2017) was run for every partition, and a consensus ML tree was generated with 100 bootstrap replicates in IQ-TREE v.1.6.5 (Nguyen *et al*., 2015). Topologies and support values from both ML inference methods were highly comparable, therefore, we only present the RAxML topologies but adding also support values from the IQ-TREE analysis. In the dataset that included available spore transcriptomic data (Beaudet *et al*., 2018, Table S2), only 17 SCOs were shared among >50% of the taxa and a phylogeny was inferred merely for visualization purposes using RAxML as described above with the concatenated alignment of retrieved SCOs.

We also reconstructed phylogenies avoiding the use of concatenated alignments for the datasets including taxa in Glomeromycota, and its sister clades (Morteriellomycota and Mucoromycota). Individual gene trees were inferred using RAxML and IQ-TREE, both with 100 bootstrap replicates. A coalescent-based species tree was inferred with ASTRAL-II using multi-locus bootstrapping (Mirarab and Warnow, 2015). Furthermore, a Bayesian phylogeny was inferred this dataset using Phylobayes (Lartillot, Lepage and Blanquart, 2009), under the site-heterogeneous CAT+GTR+G4 model on a total alignment of 144,177 amino acids. Two chains were run of ∼200k and ∼120k generations respectively and full convergence was achieved. For in-depth analysis of topologies within Glomeromycota, three additional analyses were performed. A dataset containing all Glomeromycota taxa was used to produce a splits network in IQ-TREE v.1.6.5 (Nguyen *et al*., 2015) with the command iqtree --net, the tree was then visualized in SplitsTree5 (Huson and Bryant, 2006) with a maximum dimension of 2. To increase robustness, further analyses were performed using a set of only 15 selected taxa, representing the assemblies with highest N50 and BUSCO-estimated completeness (Table S3-4), which resulted in a comprehensive dataset of 799 SCOs shared among all taxa (as opposed to 31 SCOs when including all Glomeromycota taxa). In order to visualize all topologies branching over the tree landscape, the previously inferred individual gene trees from RAxML and IQ-TREE were used to analyze the spectra of topologies with Densitree v.2.01. Finally, we used TWISST (Martin and Van Belleghem, 2017) as a topology weighting method to quantify the phylogenetic relationships between the Glomeromycota families. The method is designed for population genomic data and was thus adapted for our analysis. We used the individual gene trees from SCOs and their bootstrap values to assess the range of different topologies supported for each SCO. For visualization purposes, the concatenated set of SCOs was input as an artificial chromosome, across which we visualized the dominant topology, of three possible topologies, for each SCO.

## Results

### Presenting 21 *de novo* genome assemblies of AM fungi

Using our novel workflow for *de novo* assembly of genomes by combining single nuclei sequence data (Montoliu-Nerin *et al*., 2020), we attempted to sequence 31 AM fungal isolates, representing eight families with at least two taxa from each of 15 genera (Table S1). Spores from all 31 isolates were extracted for nuclei sorting and DNA amplification. For two of the taxa, *Ar. trappei* and *E. infrequence*, we failed to sort nuclei, and these were thus omitted from the method pipeline. After whole genome amplification with MDA on the sorted nuclei, samples from the remaining 29 isolates were screened for presence of DNA of fungal and bacterial origin by PCR amplification of rDNA sequences. Fungal DNA was successfully amplified from 25 of the isolates, while samples from four isolates did not and *Se. constrictum, Gl. microaggregatum, Gl. gold*, and a second isolate of *E. infrequence* were thus excluded from sequencing (Table S1).

After Illumina sequencing of separate amplified nuclei from the AM fungal isolates we generated *de novo* genome assemblies following the method presented as assembly workflow 3 in Montoliu-Nerin *et al*., (2020), where all reads were combined for each strain, normalized and then assembled using SPAdes. Genome assemblies ranged from 50 to 493 Mb in size, with numbers of gene predictions ranging from 11,400 to 46,500 (Table S3).

Based on the comparison of *Rh. irregularis* DAOM197198 genome assemblies, we found that single nuclei MDA and sequencing were highly accurate and efficient in our workflow. On average, around 99% of the reads mapped to both our *de novo* genome assembly and the published reference genome v.2.0 for *Rh. irregularis* DAOM197198 (Table S5). Reads from individual nuclei covered on average 50% of both assemblies (Table S5). Together these results demonstrate that reads from single amplified and sequenced nuclei fully match the published reference and that the whole genome is represented among the reads. Pair-wise alignment of the 271 BUSCO genes retrieved in both assemblies demonstrate high consistency with an average similarity of 99.7% across nucleotide alignments. Of the 271 pairwise aligned BUSCO genes, a total of 260 were identical between the two assemblies, corresponding to 96% of the retrieved BUSCO genes. However, only 60% similarity was detected in one of the 271 pairwise alignments, and ten alignments ranged in similarity between 84 and 99% (Supplementary datafile 2). High similarity in pairwise alignments of BUSCO genes retrieved from the two assemblies demonstrates that random errors possibly introduced during MDA are not retained to a large extent in the assembled genome when reads from single nuclei are combined and normalized before assembling with SPAdes. In our assembly of *Rh. irregularis* DAOM197198, 23,258 genes were predicted (Table S3) compared to 26,183 genes predicted in the published assembly of *Rh. irregularis* DAOM197198 v.2.0. (Chen *et al*., 2018). We demonstrate that our *de novo* genome assembly for *Rh. irregularis* DAOM197198 contained a largely overlapping set of genes in orthogroups present in the published *Rh. irregularis* DAOM197198 reference genome v.2.0. Across the two genome assemblies of the same strain, a total of 13,908 orthogroups were identified including 88% of all predicted genes across the two assemblies, of these, 94% were shared between the two genome assemblies (Table S6). Interestingly, both genome assemblies contain orthogroups not recovered in the other, 403 unique to v.2.0 and 380 unique to our *de novo* assembly (Table S6).

### Isolate identification in rDNA-based phylogeny

The complete rDNA operon, including SSU, ITS1, 5,8s, ITS2, and LSU regions was extracted from the 25 newly generated genome assemblies. To confirm genus level identity of the 25 isolates for which we generated genome assemblies in this study, the SSU rDNA region was extracted and placed into the Glomeromycota phylogeny of Krüger *et al*., 2012 (Fig. S1). The phylogenetic placement revealed that five isolates were originally misidentified. Four of these were removed and one re-identified for further analysis. First, the isolate *Rh. intraradices* FL208A clustered within the genus *Funneliformis* (Fig. S1), more specifically, as *F. mosseae*. Morphological examination of this strain was consistent with its original identification as *Rh. intraradices*. Because we could not verify that spores with the correct morphology had been extracted, the genome assembly was excluded from further analyses. The isolate *F. caledonius* UK203 was also identified as *F. mosseae* (Fig. S1), but kept for further analyses, since the genus placement remained correct. The isolate *Diversispora epigaea* AZ150B was phylogenetically misplaced based on its rDNA SSU sequence (Fig. S1), and the assembly had higher GC content than the rest of isolates (Table S3), probably due to bacterial contamination. We thus decided to discard this assembly since the publicly available genome assembly of *Di. epigaea* (Sun *et al*., 2018) was included in the analyses. Finally, the isolates *Ar. scheckii* CL383 and *Se. viscosum* MD215 were placed in the Paraglomeraceae family (Fig. S1), and subsequently eliminated from further analyses, as two *Paraglomus* isolates were already included. After this confirmation step, novel genome assemblies representing 21 isolates were kept for the phylogenomic analyses. A phylogenetic analysis of the entire extracted rDNA operon of the 21 isolates (Fig. S2) confirmed the expected topology found in previous analysis based on rDNA genes (Redecker *et al*., 2013).

### Phylogenomic analysis of Glomeromycota

Single copy orthologs shared among the 21 *de novo* genome assemblies, previously published genome assemblies of AM fungi and selected outgroup taxa were identified from the predicted genes and transcriptomes (Table S2-3). Published transcriptomic data from ten genera of AM fungi represented by one species each (Beaudet *et al*., 2018) is consistently placed to genus level in a phylogenetic analysis with genomic data from this and previous studies (Fig. S3). However, when combining genomic data with transcriptomic data, only 17 SCOs were shared among 50% of the taxa, thus and published trancriptomic data was thus excluded from further analysis without losing taxonomic breadth, but allowing us to work with a more comprehensive set of genes.

To place Glomeromycota in relation to its sister phyla, phylogenetic trees (Fig. S4-5) were built using 178 SCOs identified from the gene annotations and shared among 50% of the taxa included. The concatenated alignment had a length of 76,737 amino acids. Glomeromycota forms a well-supported monophyletic clade (100% bootstrap support), but the relationships between Morteriellomycota and Mucoromycota are not well resolved. Morteriellomycota is recovered as a sister clade of Glomeromycota, with bootstrap supports of 80% in the RaxML phylogeny (Fig. S5a). In the IQ-TREE phylogeny on the other hand, Morteriellomycota and Mucoromycota form a well-supported sister lineage to Glomeromycota (Fig. S5b). Their internal relation, however, is not well-supported in this analysis with only 43% bootstrap support for the separation of Morteriellomycota and Mucoromycota (Fig. S5b). The relationships among the three sister phyla thus remains unresolved in our analysis.

For in depth phylogenomic analysis of Glomeromycota, we included representatives from Morteriellomycota as sister clade, and the Mucoromycota as outgroup. A concatenated alignment of 371 SCOs shared among at least 50% of the taxa, gave a total alignment of 144,177 amino acids. All represented Glomeromycota families form well-supported monophyletic lineages in the consensus species tree (Fig. 1, Fig. S6), strongly supporting available phylogenetic inferences based on a combination of morphology and rDNA gene phylogenies (Redecker *et al*. 2013). We found, however, that the order Glomerales is polyphyletic, with taxa in Glomeraceae recovered as a sister clade to the order Diversisporales, while the family Claroideoglomeraceae forms a basal sister clade to the two, with a bootstrap support of 100% (Fig. 1, S6). In an ASTRAL reconstruction based on 371 ML individual gene trees, we observe a low multi-bootstrapping support for the node that includes Glomeraceae and Diversisporales, with a bootstrap support of 81% and 94% when using individual trees inferred with RAxML and IQ-TREE, respectively (Fig. S7).

**Figure 1.**
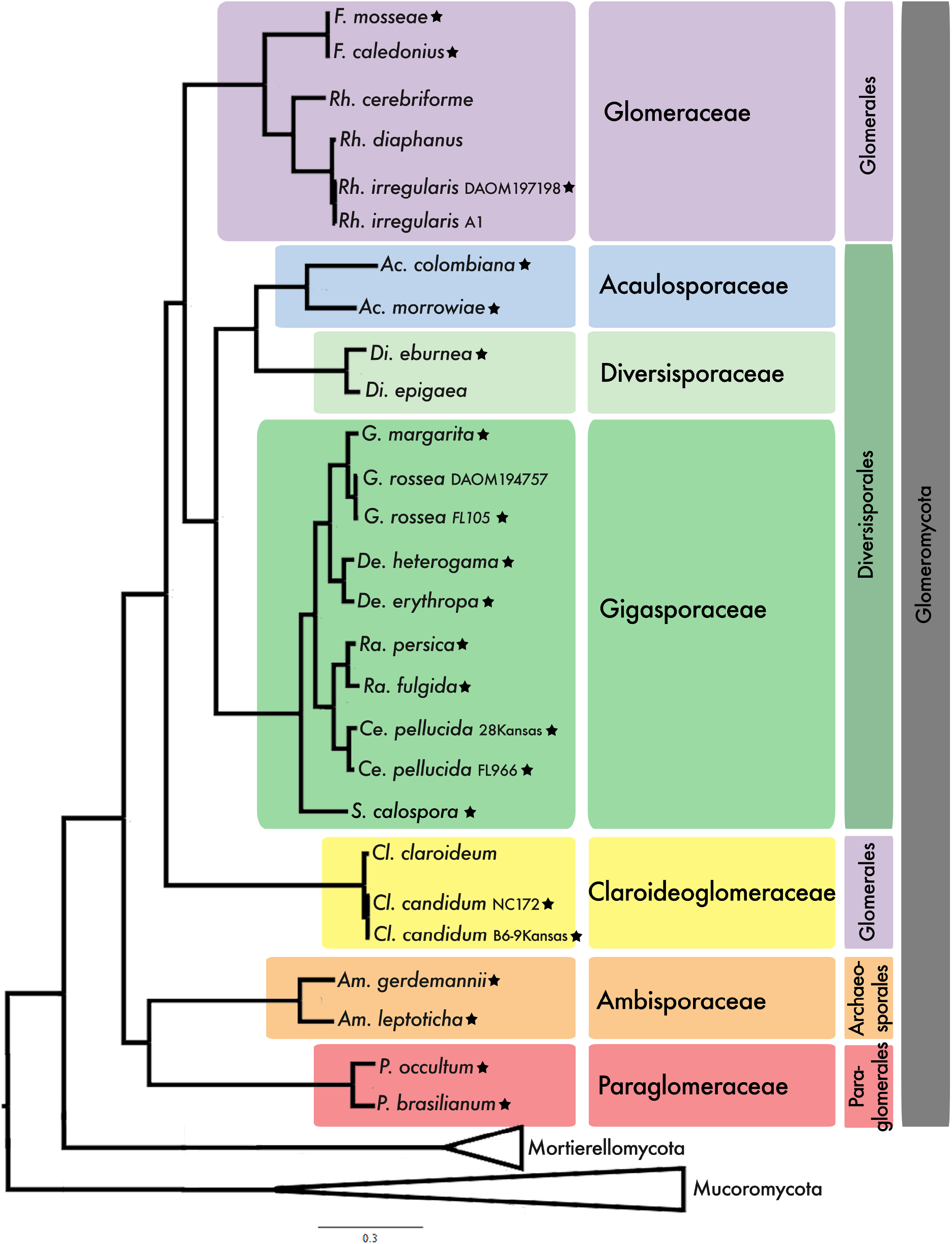
Best maximum likelihood tree inferred with RAxML from a concatenated alignment of 371 single copy orthologs shared among at least 50% of the taxa. The same topology was recovered using IQ-TREE and Bayesian inference. All nodes have a bootstrap support value of 100 in both analyses, and posterior probabilities of 1. Mucoromycota was used as outgroup. Stars following the taxon name mark newly sequenced strains from this study. Current taxonomic assignment based on Redecker *et al*., 2013 is color coded, at the levels of family and order. Strain identifers are included in the taxa label when more than one node has the same species name. See expanded tree in Fig. S6.

### Exploring conflicting topologies

To study the relationships within Glomeromycota in more detail, different datasets were produced to visualize the conflicting topologies. For this, we included only taxa within Glomeromycota to generate three datasets. First, a set of 31 SCOs shared among all Glomeromycota taxa included in this study, was concatenated to represent a total alignment of 15,443 amino acids. A second set of 1,737 SCOs represented within at least 50% of all Glomeromycota taxa included in this study (Table S2-S3), represented a total alignment of 702,801 amino acids. Finally, we produced a dataset with a reduced selection of taxa, covering all families, by picking the 15 *de novo* assembled Glomeromycota genomes with the highest quality (Table S3-4). This last dataset was used to obtain a greater number of SCOs shared among all analyzed taxa, with a total of 799 SCOs shared among all 15 represented taxa, resulting in a total alignment of 476,329 amino acids.

The main topology of the species trees (Fig. 1, S6) was not recovered in the dataset with 31 SCOs shared among all Glomeromycota (Fig. S8) where instead an alternative topology with Claroideoglomeraceae as a sister group of Diversisporales was inferred. This topology however had low bootstrap support at 52% and 59%, in RAxML and IQ-TREE respectively (Fig. S8). However, both the best ML and ASTRAL reconstruction trees, recovered the main topology (Fig. 1), when using the two other Glomeromycota datasets with the highest number of SCOs included, (Fig. S9-S12). Further analyses were performed using these more comprehensive datasets, with 1,737 SCOs shared among 50% of the Glomeromycota taxa, and 799 SCOs shared among all 15 selected Glomeromycota taxa. A Phylogenomic network reviles a clear reticulation at the base of the tree, leaving the early evolutionary relationships unresolved (Fig. 2a, S13-S14). The three most commonly observed topologies were visualized in one image using Densitree (Bouckaert and Heled, 2014), in which all trees are stacked on top of each other (Fig. 2b, S15). We find the same three topologies when using topology weighting by iterative sampling of sub-trees with TWISST (Martin and Van Belleghem, 2017, Fig. 2c, S16). This analysis shows that most genes support a predominant topology, in which we recovered Glomeraceae as sister of Diversisporales, followed by a second topology in which Glomerales is recovered as a monophyletic clade, and a third topology, in which Claroideoglomeraceae is inferred as a sister group of Diversisporales (Fig. 2b-2c, S15-S16).

**Figure 2.**
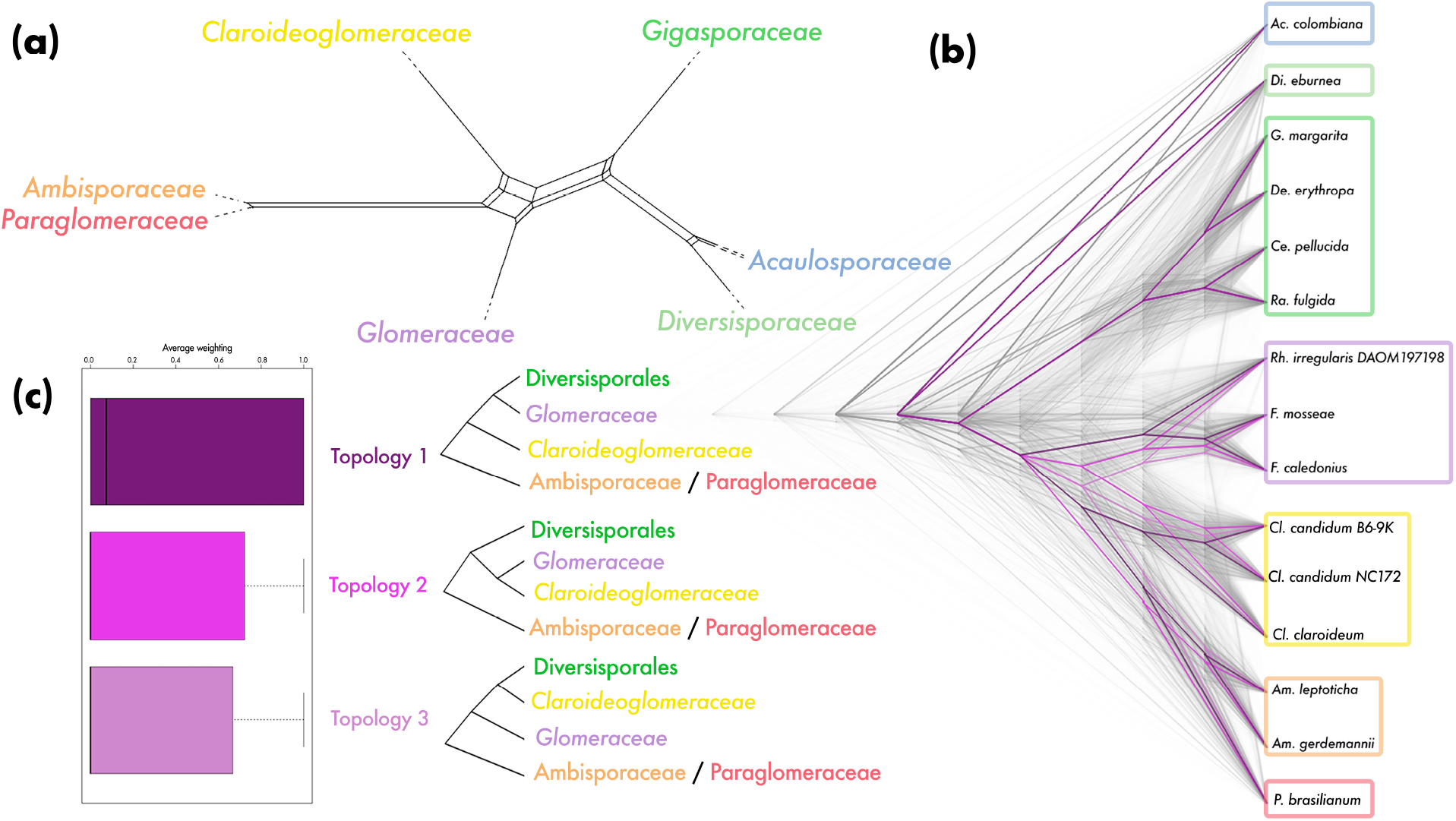
Exploring the diversity of topologies within Glomeromycota. **(a)**. IQ-TREE network analysis visualized in SplitsTree5 with maximum dimension splits filter of 2, using the dataset containing all Glomeromycota taxa, and 1,737 SCOs shared among, at least 50% of the taxa. See Fig. S13 for expanded network with branch lengths. **(b)**. Densitree of Glomeromycota, formed from stacking of 799 individual gene trees and their corresponding bootrstrapping trees for each gene tree, based on selection of 15 taxa. Three main topologies are colored in three different hues of purple, from the most common being the darkest, to the least common the lightest. For a better visualization of topology 3, the order of taxa has been rearranged in Fig. S15. **(c)**. TWISST analysis of 15 taxa of Glomeromycota, grouped in four monophyletic lineages representing the order Diversisporales, and families Glomeraceae, Claroideoglomeraceae, with Ambisporaceae / Paraglomeraceae as an outgroup. The analysis produces a topology average weighting of the three possible topologies (same colors as in (b)), using the 799 individual gene trees and their corresponding bootstrapping trees. Expanded TWISST analysis in S16.

## Discussion

Glomeromycota encompass all known AM fungi with their characteristic life cycle involving an obligate association with plants (Bonfante & Venice 2020) as well as the exceptional fungal taxa *Geosiphon pyriformis* which forms a mutualistic symbiosis with the cyanobacteria *Nostoc punctiforme* (Malar *et al*., 2021). In the current study we present a four-fold increase in the number of AM fungal genomes available, which was achieved thanks to the development of a workflow for genome assembly from multiple individually amplified and sequenced nuclei (Montoliu-Nerin *et al*., 2020).

The current workflow for generating *de novo* reference genomes of AM fungi was developed by our team to circumvent the need for culturing AM fungi for genomic studies (Montoliu-Nerin *et al*., 2020). Read mapping of data from 24 individually amplified and sequenced *Rh. irregularis* DAOM197198 nuclei demonstrates near complete coverage of the published *Rh. irregularis* DAOM197198 v.2.0 reference genome (Table S5), suggesting that separate amplification of multiple nuclei compensates for uneven amplification of individual nuclei. Consistent recovery of orthogroups in our *de novo* genome assembly of *Rh. irregularis* DAOM197198 (Table S6) and evidence that mostly identical BUSCO genes are recovered from both assemblies provides further support that the presented workflow generates gene sequence data suitable for phylogenomic analysis. We anticipate that the release of these novel genome assemblies will become an important resource for the future study of AM fungi, supplementing the previous AM fungal genomes (Tisserant *et al*., 2013; Lin *et al*., 2014; Chen *et al*., 2018; Kobayashi *et al*., 2018; Sun *et al*., 2018; Morin *et al*., 2019; Montoliu-Nerin *et al*., 2020).

Based on the most comprehensive taxon sampling thus far we present a well-supported species tree for AM fungi. We find that the seven family level lineages included in the analysis represent well supported monophyletic lineages. While the order Diversisporales, including three families, was recovered as monophyletic we found that the order Glomerales with the two families Glomeraceae and Claroideoglomeraceae was not. Comprehensive phylogenetic studies with wide taxon sampling representing AM fungi have thus far mostly used rDNA sequences from trap cultures (Redecker *et al*., 2013) and environmental samples (Krüger *et al*., 2012) and recover Glomerales as monophyletic based on these markers. Similarly, our phylogenetic reconstructions using only the extracted rDNA operon from the *de novo* assembled genomes support Glomerales as monophyletic (Fig S2). Glomerales was previously found to be polyphyletic in a phylogenomic analyses using spore transcriptomic data from nine AM fungal species, where *Claroideoglomus* was recovered as a well-supported clade with Ambisporaceae and Paraglomeraceae (Beaudet *et al*., 2018). In contrast to Beaudet *et al*. (2018), we recovered Glomeraceae as a sister group of Diversisporales with Claroideoglomeraceae being the basal sister group of the two (Fig. 1-2). Previous phylogenomic studies using whole genomic data had not yet observed this topology due to limited taxon sampling (Morin *et al*., 2019). Further, the relation of Glomeromycota to its to closest sister lineages, Mucoromycota and Morteriellomycota, have not yet been resolved with strong support, and based on available data the relation is best described as a polytomy (Li *et al*., 2021). Interestingly, with the addition of a considerable number of AM fungal genomes presented in this study, we recover yet a new topology among the three sister lineages (Fig. S4-5). This further highlights the need for increased taxon sampling in the sister lineages, particularly in Morteriellomycota.

The placement of Glomeraceae as a sister group of Diversisporales is well supported in the species tree (Fig. 1), but alternative topologies were indicated by the lower support based on coalescence methods (Fig. S7). A more detailed analysis of the different topologies was possible based on the single gene trees from orthologs shared among the included Glomeromycota taxa, after removing taxa belonging to Morteriellomycota and Mucoromycota, in order to have a larger pool of SCOs (Fig. 2, S8-S16). The most common topology groups Glomeraceae and Diversisporales, as in the species tree; a second one, which recovers Glomerales as monophyletic; and a third places Claroideoglomeraceae as a sister group of Diversisporales (Fig. 2, S15-S16).

It is possible that the tree topology discordances are due to incomplete lineage sorting (Maddison and Knowles, 2006), caused by long coalescence times which complicates the assessment of an accurate evolutionary history. Different topologies could also result from gene flow among AM fungal lineages. Documented gene family expansions correlated with genome size in AM fungi (Tang *et al*., 2016), could distort phylogenetic histories since gene expansions and contractions can cause misidentification of SCOs, resulting in alignments between paralogs present as single copy with different evolutionary origins and histories. A better understanding on how variation in gene content and copy number variation influenced the different topologies could be achieved with a deeper phylogenetic study into the whole repertoire of paralogs, moving one step further from SCOs, which would also allow us to look more closely into the possible correlation between gene function and the different evolutionary histories.

## Conclusion

In the current study we present a considerable increase in the number of AM fungal genome assemblies available, thanks to the development of single nuclei sequencing and *de novo* assembling in AM fungi. We conclude that our *de novo* genome assemblies provide a satisfactory representation of the genome content. While the assemblies generated with our workflow are fragmented, we demonstrate that genome content is well recovered across nuclei, despite variation in sequencing depth due to MDA. Further, we demonstrate that gene content is soundly recovered in the *de novo* genome assemblies. We present a phylogenomic analysis of AM fungi based on the most comprehensive taxon sampling across Glomeromycota to date. Our results support current family-level classification and in the first and most broadly supported tree topology, the order Glomerales is polyphyletic, with the family Glomeraceae being recovered as a sister group to the order Diversisporales, while Claroideoglomeraceae is recovered as an outgroup of the two. The new genome data presented cover seven families of the phylum Glomeromycota and are expected to be a valuable contribution to the AM fungal research community.

## Supporting information

Annex I

Annex II

## Acknowledgments

Funding for this project was provided by ERC (678792). We thank Y. Strid for assistance in the lab and J. Morton and W. Wheeler at INVAM culture collection. Nuclei sorting and whole genome amplification was done at the SciLifeLab Microbial Single Cell Genomics Facility at Uppsala University. Sequencing was performed by the SNP&SEQ Technology Platform at NGI Sweden and SciLife Laboratory, Uppsala, supported by the VR and the KAW. Computational analyses were performed on resources provided by SNIC through UPPMAX. VK was financially supported by the Knut and Alice Wallenberg Foundation as part of the National Bioinformatics Infrastructure Sweden at SciLifeLab. Open access funding provided by Uppsala University. The paper was greatly improved thank to comments from three anonymous reviewers.

## Author contribution

M.M.N. initiated the project together with A.R. and J.D.B. and developed the analysis together with M.S.G. and H.J.. C.B. did the nuclei sorting and whole genome amplification. M.M.N. performed the bioinformatic analyses together with M.S.G., V.K. developed the annotation workflow which was ran by M.M.N and M.S.G.. M.M.N. wrote the manuscript with A.R. and H.J., with input from all the authors.

