## Supplementary Information for "In-depth phylogenomic analysis of arbuscular mycorrhizal fungi based on a comprehensive set of de novo genome assemblies"

**Supplementary tables**

**Table S1.** Table of all isolates attempted for sequencing.

**Table S2.** Published genomic data used in phylogenomic analysis.

**Table S3.** Whole genome assembly information for all *de novo* sequenced isolates.

**Table S4.** Complete BUSCO statistics for all *de novo* sequenced isolates

**Table S5**. Assembly coverage of reads mapped to two assemblies of *Rh. irregularis* DAOM197198

**Table S6.** Summary statistics from OrthoFinder across two assemblies of *Rh. irregularis* DAOM197198

**Supplementary figures**

**Figure S1.** SSU rDNA phylogeny to confirm isolate identity.

**Figure S2.** Phylogenetic analysis of complete rDNA operon of 21 isolates this study.

**Figure S3.** Best ML tree including transcriptomic data from Beaudet *et al.*, 2018.

**Figure S4.** Visualization of best ML tree including Glomeromycota and its sister lineages.

**Figure S5.** Expanded version of ML tree in Fig. S4 a) RAxML and b) IQ-TREE

**Figure S6.** Expanded version of ML tree in Fig. 1

**Figures S7.** Multi-locus bootstrapping ASTRAL tree from 371 individual gene trees

**Figure S8**. Best ML tree of Glomeromycota from 31 single copy orthologs shared among all taxa.

**Figure S9.** Best ML tree of Glomeromycota, 1,737 SCOs shared among, at least, 50% of taxa

**Figure S10.** Multi-locus bootstrapping ASTRAL tree of Glomeromycota, from 1,737 individual gene trees

**Figure S11.** Best ML tree of Glomeromycota, 799 SCOs shared among 15 selected taxa.

**Figure S12.** Multi-locus bootstrapping ASTRAL tree of Glomeromycota, from 799 individual gene trees

**Figure S13.** Expanded network from Figure 2a,

**Figure S14.** Network from IQ-TREE network analysis, using 799 single gene trees shared among 15 selected taxa

**Figure S15** Same Densitree as in Figure 2b based on 15 selected taxa, with order of taxa rearranged to visualize topology 3, and Densitrees with IQ-TREE single gene trees

**Figure S16.** TWISST analysis on a selection of 15 taxa

**Table S1.** Table of all isolates for which *de novo* genome sequencing was attempted. Organized according to classification on INVAM homepage (2020-08-01). The whole genome amplification (WGA) kit used for amplification of the single nuclei DNA is indicated under (WGA) for each isolate. Number of (#) spores indicates how many spores were pooled to extract nuclei for FACS, and number of nuclei shows how many nuclei were individually selected for sequencing, resulting in the same number of libraries successfully sequenced.

| **Family** | **Genus** | **Species** | **Strain** | **Collection^3^** | **Nuclei sorted^4^** | **WGA** | **# spores** | **# nuclei** |
| --- | --- | --- | --- | --- | --- | --- | --- | --- |
| **Glomeraceae** | Funneliformis | *Funneliformis mosseae* | 87-6 pot B 2015 | Kansas |  | Epicentre | ~15 | 20 |
|  |  | *Funneliformis caledonius* | UK204 | INVAM |  | Qiagen | ~15 | 24 |
|  | Septoglomus | *Septoglomus viscosum* | MD215 | INVAM |  | Qiagen | ~15 | 20 |
|  |  | *Septoglomus constrictum* | KS890 | INVAM |  |  | ~15 | - |
|  | Glomus | *Glomus microaggregatum* | UT126B | INVAM |  |  | ~15 | - |
|  |  | *Glomus gold* | KS906B | INVAM |  |  | ~15 | - |
|  | Rhizophagus | *Rhizophagus irregularis* | DAOM197198 | Canada |  | Qiagen | 1 | 24 |
|  |  | *Rhizophagus intraradices* | FL208A | INVAM |  | Epicentre | 1 | 24 |
| **Acaulosporaceae** | Acaulospora | *Entrophospora infrequence* ^1^ | 110 2015 | Kansas | No |  | ~15 | - |
|  |  | *Entrophospora infrequence* ^1^ | CA203 | INVAM |  |  | ~15 | - |
|  |  | *Acaulospora colombiana* | CL356 | INVAM |  | Qiagen | ~15 | 15 |
|  |  | *Acaulospora morrowiae* | CL551 | INVAM |  | Qiagen | ~15 | 19 |
| **Diversisporaceae** | Diversispora | *Diversispora epigaea* | AZ150B | INVAM |  | Qiagen | ~15 | 7 |
|  |  | *Diversispora eburnea* | AZ414A | INVAM |  | Qiagen | ~15 | 24 |
| **Gigasporaceae** | Gigaspora | *Gigaspora margarita* | 120-4 pot B 10/14 | Kansas |  | Qiagen | ~15 | 24 |
|  |  | *Gigaspora rosea* | FL105 | INVAM |  | Epicentre | ~15 | 20 |
|  | Dentiscutata | *Dentiscutata heterogama* | IL203A | INVAM |  | Qiagen | ~15 | 7 |
|  |  | *Dentiscutata erythropus* | MA453B | INVAM |  | Qiagen | ~15 | 24 |
|  | Cetraspora | *Cetraspora pellucida* | FL966 | INVAM |  | Qiagen | ~15 | 24 |
|  |  | *Cetraspora pellucida* ^2^ | 28 12/20/2015 | Kansas |  | Epicentre | ~15 | 17 |
|  | Racocetra | *Racocetra persica* | MA461A | INVAM |  | Qiagen | ~15 | 24 |
|  |  | *Racocetra fulgida* | IN212 | INVAM |  | Qiagen | ~15 | 22 |
|  | Scutellospora | *Scutellospora calospora* | AU212A | INVAM |  | Epicentre | ~15 | 13 |
| **Claroideoglomeraceae** | Claroideoglomus | *Claroideoglomus candidum* | NC172 | INVAM |  | Epicentre | 1 | 24 |
|  |  | *Claroideoglomus candidum* | (CCK) pot B 6-9 | Kansas |  | Qiagen | 7 | 24 |
| **Paraglomeraceae** | Paraglomus | *Paraglomus occultum* | IA702 | INVAM |  | Epicentre | ~15 | 23 |
|  |  | *Paraglomus brasilianum* | BR232B | INVAM |  | Epicentre | ~15 | 20 |
| **Archaeosporaceae** | Archaeospora | *Archaeospora trappei* | IL203B | INVAM | No |  | ~15 | - |
|  |  | *Archaeospora schencki* | CL383 | INVAM |  | Qiagen | ~15 | 24 |
| **Ambisporaceae** | Ambispora | *Ambispora leptoticha* | FL130A | INVAM |  | Qiagen | ~15 | 22 |
|  |  | *Ambispora gerdemannii* | MT106 | INVAM |  | Qiagen | ~15 | 24 |

1. Taxonomic placement of *Entrophspora infrequences* remains unresolved
2. The strain was previously named *Scutellospora pellucida*. It is the same culture as INVAM collection *Cetraspora pellucida* (INVAM IN211)
3. Collections from where the strains were obtained: Kansas: James D. Bever’s lab, University of Kansas, USA; INVAM: International culture collection of (vesicular) arbuscular mycorrhizal fungi (INVAM) at West Virginia University, Morgantown, WV, USA; Canada: Agriculture and Agri-food Canada, Government of Canada.
4. Shadowed boxes indicate successful nuclei sorting.

**Table S2.** Published data used in phylogenomic analyses. Annotations of the whole genome assemblies were downloaded from their original source. Final column shows in which figures each isolate was included.

| **Phyla** | **Species** | **Isolate** | **Publication** | **Used for Fig.** |
| --- | --- | --- | --- | --- |
| Glomeromycota | *Rhizophagus irregularis* | A1 | Chen *et al.*, 2018 | 1-2,  S3-S10, S13 |
|  | *Rhizophagus diaphanus* | MUCL43196 | Morin *et al.*, 2019 | 1-2,  S3-S10, S13 |
|  | *Rhizophagus cerebriforme* | DAOM227022 | Morin *et al.*, 2019 | 1-2,  S3-S10, S13 |
|  | *Rhizophagus irregularis* | DAOM234181 | Beaudet *et al.*, 2018 | S3 |
|  | *Funneliformis mosseae* | DAOM236685 | Beaudet *et al.*, 2018 | S3 |
|  | *Acaulospora morrowiae* | CR315B | Beaudet *et al.*, 2018 | S3 |
|  | *Diversispora epigaea* | IT104 | Sun *et al.*, 2018 | 1-2,  S3-S10, S13 |
|  | *Diversispora versiforme* | W475-40 | Beaudet *et al.*, 2018 | S3 |
|  | *Gigaspora rosea* | DAOM194757 | Morin *et al.*, 2019 | 1-2,  S3-S10, S13 |
|  | *Racocetra castanea* | BEG 1 | Beaudet *et al.*, 2018 | S3 |
|  | *Scutellospora calospora* | IL209 | Beaudet *et al.*, 2018 | S3 |
|  | *Claroideoglomus claroideum* | DAOM234280 | Beaudet *et al.*, 2018 | S3 |
|  | *Claroideoglomus claroideum* | SA101 | Montoliu-Nerin *et al.*, 2020 | 1-3,  S3-S16 |
|  | *Ambispora lepototicha* | JA116 | Beaudet *et al.*, 2018 | S3 |
|  | *Paraglomus brasilianum* | DAOM240472 | Beaudet *et al.*, 2018 | S3 |
| Morteriellomycota | *Morteriella elongata* | AG 77 | Uehling *et al.*, 2017 | 1, S3-S7 |
|  | *Lobosporangium transversale* | NRR 3116 | Mondo *et al.*, 2017 | 1, S3-S7 |
| Mucoromycota | *Endogone sp.* | FLAS 59071 | Chang *et al.*, 2019 | 1, S3-S7 |
|  | *Jimgerdemannia lactiflua* | OSC 166217 | Chang *et al.*, 2019 | 1, S3-S7 |
|  | *Jimgerdemannia flammicorona* | AD 002 | Chang *et al.*, 2019 | 1, S3-S7 |
|  | *Syncephalastrum racemosum* | NRRL 2496 | Mondo *et al.*, 2017 | 1, S3-S7 |
|  | *Lichtheimia corymbifera* | JMRC FSU 9682 | Schwartze *et al.*, 2014 | 1, S3-S7 |
|  | *Hesseltinella vesiculosa* | NRRL 3301 | Mondo *et al.*, 2017 | 1, S3-S7 |
|  | *Absidia repens* | NRRL 1336 | Mondo *et al.*, 2017 | 1, S3-S7 |
|  | *Saksenaea vasiformis* | B4078 | Chibucos *et al.*, 2016 | 1, S3-S7 |
|  | *Phycomyces blakesleeanus* | NRRL 1555 | Corrochano *et al.*, 2016 | 1, S3-S7 |
|  | *Rhizopus microsporus var. microsporus* | ATCC 52813 | Mondo, Lastovetsky, *et al.*, 2017 | 1, S3-S7 |
|  | *Rhizopus microsporus var. chinensis* | CCTCCM 201021 | Wang *et al.*, 2013 | 1, S3-S7 |
|  | *Mucor circinelloides* | CBS 277 49 | Corrochano *et al.*, 2016 | 1, S3-S7 |
|  | *Rhizopus delemar* | RA99 880 | Ma *et al.*, 2009 | 1, S3-S7 |
| Basidiomycota | *Laccaria bicolor* | S238N-H82 | Martin *et al.*, 2008 | 1  S4-S5 |
|  | *Ustilago maydis* | 521, DSMZ14603 | Kämper *et al.*, 2006 | 1,  S4-S5 |
|  | *Puccinia striiformis f. sp. tritici* | 104 E137 A- | Schwessinger *et al.*, 2018 | 1,  S4-S5 |
| Ascomycota | *Tuber melanosporum* | Mel28 | Martin *et al.*, 2010 | 1,  S4-S5 |
|  | *Schizosaccharomyces pombe* | - | Wood *et al.*, 2002 | 1,  S4-S5 |
|  | *Yarrowia lypolitica* | FKP355 | Pomraning *et al.*, 2018 | 1,  S4-S5 |

**Table S3**. Whole genome assembly information for all *de novo* genome assemblies generated in this study. Showing results from the QUAST analysis (Estimated size, Size, #Contigs, N50, Largest contig and GC), estimated completeness analysis (BUSCO), results from gene prediction and repeat annotation pipeline (# Genes and (content)), Repeat content). Final column indicates in which figures each taxa was included.

| **Species** | **Estimated size**  **(Mb)** | **Size**  **(Mb)** | **# Contigs** | **N50** | **Largest contig**  **(Kb)** | **GC**  **(%)** | **BUSCO**  **(%)** | **# Genes and (content,**  **Mb)** | **Repeat content**  **(Mb)** | **Used for Fig.** |
| --- | --- | --- | --- | --- | --- | --- | --- | --- | --- | --- |
| *F. mosseae* | 156.54 | 145.72 | 27,134 | 11,988 | 97.04 | 25.22 | C: 86, F: 6 | 16,857 (42.65) | 77.14 | 1-2, S1-S16 |
| *F. caledonius* | 150.38 | 146.16 | 31,052 | 9,933 | 81.73 | 26.00 | C: 89, F: 4 | 17,946 (41.00) | 75.48 | 1-2, S1-S16 |
| *Se. viscosum* | 86.48 | 69.04 | 13,413 | 14,169 | 159.44 | 36.25 | C: 90, F: 4 | 14,815 (37.81) | 17.27 | S1 |
| *Rh. irregularis* DAOM197198 | 133.86 | 116.45 | 15,939 | 21,163 | 222.15 | 27.28 | C: 94, F: 1 | 23,258 (68.13) | 33.80 | 1-2, S1-S16 |
| *Rh. intraradices* | 133.86 | 116.45 | 15,939 | 21,163 | 222.15 | 27.28 | C: 94, F: 1 | 23,258 (68.13) | 33.80 | S1 |
| *Ac. colombiana* | 247.44 | 299.81 | 7,5131 | 8,208 | 86.29 | 29.39 | C: 78, F: 9 | 14,505 (23.78) | 240.52 | 1-2, S1-S16 |
| *Ac. morrowiae* | 154.35 | 218.77 | 64,585 | 6,189 | 54.03 | 27.89 | C: 74, F: 14 | 18,394 (31.29) | 159.05 | 1-2, S1-S10, S13 |
| *Di. eburnea* | 87.85 | 65.11 | 9,803 | 22,612 | 184.86 | 25.84 | C: 92, F: 3 | 12,017 (37.55) | 25.62 | 1-2, S1-S16 |
| *Di. epigaea* | 54.32 | 64.27 | 16,527 | 8,159 | 138.06 | 38.11 | C: 76, F: 12 | 15,972 (29.58) | 10.12 | S1 |
| *G. margarita* | 492.60 | 574.57 | 173,647 | 5,875 | 76.23 | 27.40 | C: 88, F: 3 | 46,492 (82.99) | 378.98 | 1-2, S1-S16 |
| *G. rosea* | 200.66 | 240.31 | 90,143 | 4,479 | 50.41 | 28.74 | C: 46, F: 13 | 26,343 (36.22) | 154.42 | 1-2, S1-S10, S13 |
| *De. heterogama* | 167.37 | 181.57 | 61,528 | 4,991 | 40.56 | 27.81 | C: 44, F: 11 | 16,277 (23.62) | 121.82 | 1-2, S1-S10, S13 |
| *De. erythropus* | 248.76 | 293.54 | 73,964 | 7,545 | 61.80 | 28.34 | C: 87, F: 4 | 28,764 (58.16) | 190.17 | 1-2, S1-S16 |
| *Ra. persica* | 176.70 | 353.07 | 181,541 | 2,886 | 30.66 | 27.09 | C: 47, F: 24 | 37,045 (36.01) | 238.55 | 1-2, S1-S10, S13 |
| *Ra. fulgida* | 212.27 | 295.99 | 105,307 | 4,839 | 74.97 | 28.40 | C: 68, F: 14 | 19,906 (29.68) | 205.34 | 1-2, S1-S16 |
| *Ce. pellucida 28 K*^1^ | 214.21 | 226.81 | 79,972 | 4,932 | 51.05 | 26.61 | C: 43, F: 12 | 18,058 (26.55) | 164.40 | 1-2, S1-S10, S13 |
| *Ce. pellucida FL966* | 418.72 | 435.08 | 90,930 | 10,017 | 77.05 | 25.97 | C: 89, F: 5 | 22,053 (51.29) | 325.42 | 1-2, S1-S16 |
| *S. callospora* | 143.92 | 147.04 | 50,548 | 4,999 | 47.38 | 26.36 | C: 29, F: 15 | 11,479 (16.87) | 107.94 | 1-2, S1-S10, S13 |
| *Cl. candidum* NC172 | 86.47 | 68.12 | 12,232 | 15,216 | 101.84 | 27.83 | C: 88, F: 3 | 15,761 (42.66) | 21.83 | 1-2, S1-S16 |
| *Cl. candidum B6-9K* | 87.88 | 69.90 | 12,603 | 15,877 | 114.29 | 27.86 | C: 87, F: 4 | 16,088 (43.88) | 22.68 | 1-2, S1-S16 |
| *Ar. schenckii* | 86.41 | 89.56 | 17,380 | 12,048 | 123.55 | 36.28 | C: 86, F: 8 | 19,726 (47.95) | 24.04 | S1 |
| *Am. gerdemannii* | 102.08 | 87.97 | 19,363 | 10,266 | 96.67 | 28.22 | C: 90, F: 3 | 13,690 (33.64) | 42.05 | 1-2, S1-S16 |
| *Am. leptoticha* | 163.75 | 197.20 | 58,506 | 5,886 | 99.48 | 23.62 | C: 90, F: 3 | 14,642 (30.94) | 144.29 | 1-2, S1-S16 |
| *P. occultum* | 49.53 | 50.06 | 8,053 | 16,033 | 146.03 | 36.55 | C: 76, F: 7 | 11,385 (30.35) | 10.57 | 1-2, S1-S10, S13 |
| *P. brasilianum* | 61.76 | 58.47 | 7,115 | 21,894 | 153.20 | 36.59 | C: 90, F: 3 | 11,842 (36.52) | 15.09 | 1-2, S1-S16 |

1. The strain was named *Scutellospora pellucida* in Bever lab collection (K for Kansas). It is the same culture as INVAM collection *Cetraspora pellucida* (INVAM IN211)

**Table S4.** Complete BUSCO statistics for all *de novo* genome assemblies generated using single nuclei sequencing and assembly of combined and normalized reads using SPAdes. Number of genes (out of a total of 290 conserved single copy genes) are listed that were retrieved in the four categories (Compete single copy, Complete duplicated, Fragmented or Missing). Column “15 selected” marks the 15 genome assemblies selected for in-depth analysis of Glomeromycota.

| **Species** | **Complete single copy** | **Complete duplicated** | **Fragmented** | **Missing** | **15 selected** |
| --- | --- | --- | --- | --- | --- |
| *F. mosseae* | 248 | 1 | 18 | 23 | * |
| *F. caledonius* | 256 | 2 | 12 | 20 | * |
| *Se. viscosum* | 257 | 5 | 11 | 17 |  |
| *Rh. irregularis* | 271 | 1 | 4 | 14 | * |
| *Rh. intraradices* | 249 | 2 | 14 | 25 |  |
| *Ac. colombiana* | 188 | 37 | 25 | 40 | * |
| *Ac. morrowiae* | 207 | 8 | 42 | 33 |  |
| *Di. eburnea* | 265 | 3 | 9 | 13 | * |
| *Di. epigaea* | 181 | 41 | 36 | 32 |  |
| *G. margarita* | 252 | 4 | 9 | 25 | * |
| *G. rosea* | 131 | 1 | 38 | 120 |  |
| *De. heterogama* | 127 | 1 | 31 | 131 |  |
| *De. erythropus* | 246 | 5 | 13 | 26 | * |
| *Ra. persica* | 135 | 0 | 70 | 85 |  |
| *Ra. fulgida* | 195 | 3 | 40 | 52 | * |
| *Ce. pellucida 28* K^1^ | 124 | 1 | 34 | 131 |  |
| *Ce. pellucida FL966* | 255 | 4 | 14 | 17 | * |
| *S. callospora* | 84 | 1 | 42 | 163 |  |
| *Cl. claroideum* |  |  |  |  | * |
| *Cl. candidum* | 222 | 33 | 10 | 25 | * |
| *Cl. candidum K*^2^ | 223 | 30 | 12 | 25 | * |
| *Ar. schenckii* | 200 | 50 | 23 | 17 |  |
| *Am. gerdemannii* | 256 | 6 | 9 | 19 | * |
| *Am. leptoticha* | 254 | 5 | 10 | 21 | * |
| *P. occultum* | 219 | 3 | 40 | 52 |  |
| *P. brasilianum* | 258 | 4 | 10 | 18 | * |

1. The strain was named *Scutellospora pellucida* in Bever lab collection (K for Kansas). It is the same culture as INVAM collection *Cetraspora pellucida* (INVAM IN211)

2. K is added to indicate that this is the *Claroideoglomus candidum* from Kansas (CCK) pot B 6-9 is from the Bever lab collection.

**Table S5**. Mapping of single nuclei reads of *Rh. irregularis* DAOM197198 to reference genome assembly v.2.0 (ref v.2.0), and the *de novo* assembly produced in this study (*de novo*). Percentage of mapped reads from each nucleus (1-24) and percentage assembly covered by the reads from 24 individually amplified and sequenced nuclei.

|  | % mapped reads | | % of assembly covered  (>= 1X) | |
| --- | --- | --- | --- | --- |
|  | ref v.2.0 (%) | *de novo* (%) | ref v.2.0 (%) | *de novo* (%) |
| Rh. irregularis - Nucleus 1 | 99.11 | 99.61 | 50.84 | 53.42 |
| Rh. irregularis - Nucleus 2 | 99.35 | 99.58 | 25.83 | 26.72 |
| Rh. irregularis - Nucleus 3 | 99.39 | 99.63 | 51.75 | 54.34 |
| Rh. irregularis - Nucleus 4 | 99.27 | 99.62 | 54.31 | 57.16 |
| Rh. irregularis - Nucleus 5 | 99.36 | 99.67 | 50.34 | 52.85 |
| Rh. irregularis - Nucleus 6 | 99.06 | 99.52 | 52.90 | 55.68 |
| Rh. irregularis - Nucleus 7 | 99.26 | 99.67 | 51.65 | 54.13 |
| Rh. irregularis - Nucleus 8 | 93.52 | 99.65 | 39.68 | 41.49 |
| Rh. irregularis - Nucleus 9 | 99.31 | 99.62 | 64.93 | 68.37 |
| Rh. irregularis - Nucleus 10 | 99.39 | 99.69 | 65.17 | 68.58 |
| Rh. irregularis - Nucleus 11 | 99.34 | 99.65 | 66.51 | 70.01 |
| Rh. irregularis - Nucleus 12 | 99.45 | 99.68 | 20.97 | 22.20 |
| Rh. irregularis - Nucleus 13 | 99.30 | 99.67 | 62.98 | 66.20 |
| Rh. irregularis - Nucleus 14 | 92.37 | 92.79 | 53.95 | 56.48 |
| Rh. irregularis - Nucleus 15 | 99.14 | 99.53 | 85.38 | 90.25 |
| Rh. irregularis - Nucleus 16 | 98.83 | 99.71 | 47.19 | 49.65 |
| Rh. irregularis - Nucleus 17 | 98.80 | 99.58 | 47.36 | 49.62 |
| Rh. irregularis - Nucleus 18 | 99.44 | 99.63 | 41.46 | 43.41 |
| Rh. irregularis - Nucleus 19 | 99.33 | 99.65 | 64.42 | 67.94 |
| Rh. irregularis - Nucleus 20 | 99.24 | 99.57 | 48.54 | 50.98 |
| Rh. irregularis - Nucleus 21 | 99.45 | 99.65 | 37.99 | 39.70 |
| Rh. irregularis - Nucleus 22 | 99.36 | 99.68 | 44.36 | 46.40 |
| Rh. irregularis - Nucleus 23 | 99.24 | 99.60 | 32.81 | 34.04 |
| Rh. irregularis - Nucleus 24 | 99.07 | 99.42 | 39.91 | 41.85 |
| **Average** | **98.72** | **99.34** | **50.05** | **52.56** |

**Table S6.** OrthoFinder overall statistics when searching for orthogroups shared (In both) between *Rh. irregularis* DAOM197198 reference genome v.2.0 (Chen et al., 2018) and the *de novo* assembly of the same strain produced in this study. Followed by numbers for the assemblies separately.

| Number of “species” (i.e. genome assemblies) | In both | v.2.0 | *de novo* |
| --- | --- | --- | --- |
| Number of genes | 49,451 | 26,183 | 23,268 |
| Number of genes in orthogroups | 43,894 | 23,429 | 20,465 |
| Number of unassigned genes | 5,557 | 2,754 | 2,803 |
| Percentage of genes in orthogroups | 88.8 | 89.5 | 88.0 |
| Percentage of unassigned genes | 11.2 | 10.5 | 12.0 |
| Number of orthogroups | 13,908 | 13,528 | 13,505 |
| Number of species-specific orthogroups | 783 | 403 | 380 |
| Number of genes in species-specific orthogroups | 4,264 | 2,638 | 1,626 |
| Percentage of genes in species-specific orthogroups | 8.6 | 10.1 | 7.0 |
| Mean orthogroup size | 3.2 |  |  |
| Median orthogroup size | 2.0 |  |  |
| G50 (assigned genes) | 3 |  |  |
| G50 (all genes) | 2 |  |  |
| O50 (assigned genes) | 3,092 |  |  |
| O50 (all genes) | 4,324 |  |  |
| Number of orthogroups with all species present | 13,125 |  |  |
| Number of single-copy orthogroups | 10,138 |  |  |

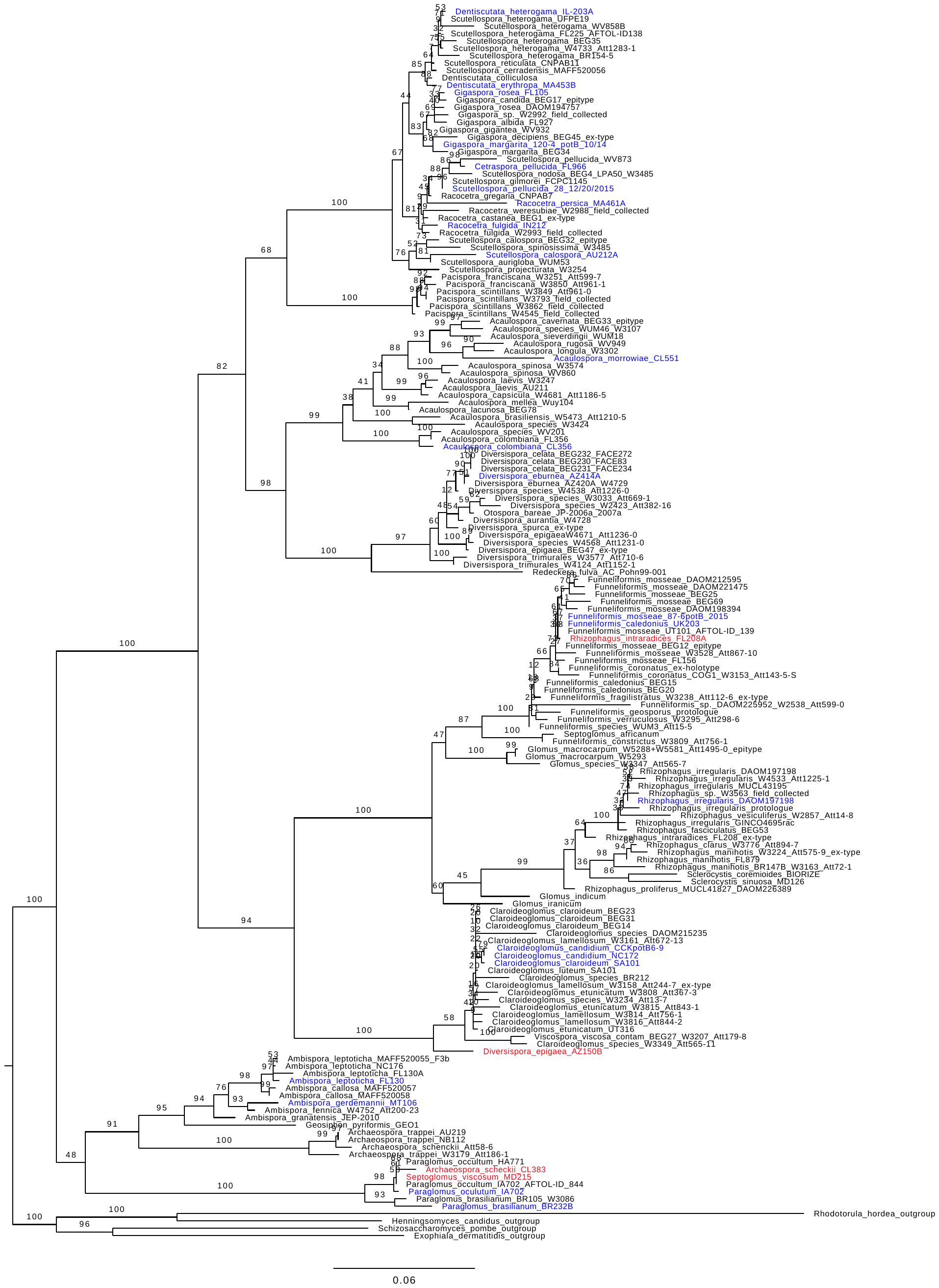
**Figure S1.** Best maximum likelihood RAxML tree from an alignment of the extracted small subunit (SSU) of the rDNA from the newly assembled genomes (in color) with SSU sequences published in Krüger *et al.* 2012 (in black). Isolates sequenced in the current study are labeled according to initial information from culture collections (Table S1). Taxon names in blue indicating that placement correspond with genus name while names in red indicate that isolate were removed from further analysis because of suspected contamination or misidentification due to inconsistent placement based on taxon name. Taxa in blue were included in downstream analysis. Branch labels show bootstrap support (100 replicates).

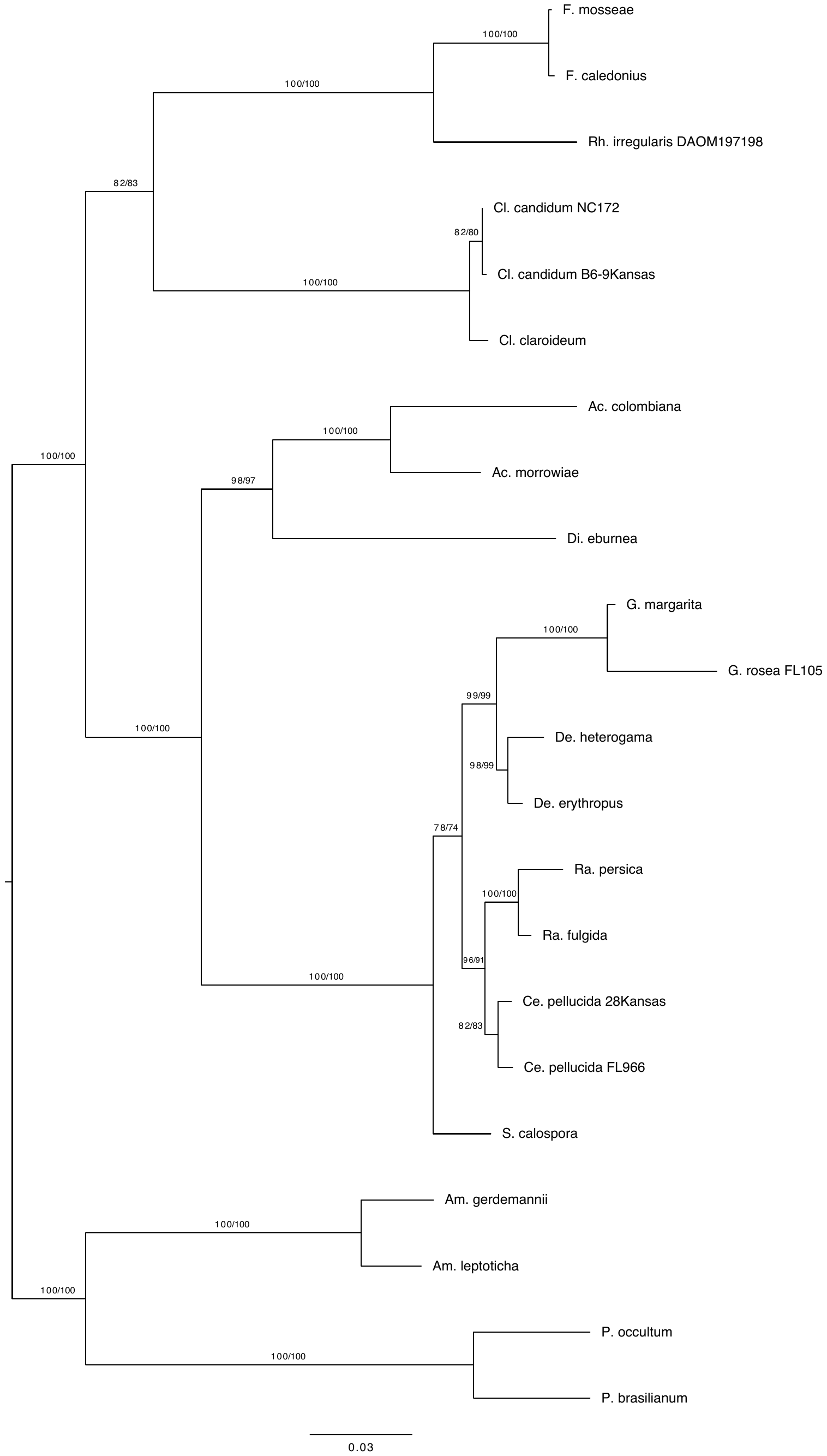

**Figure S2.** Best maximum likelihood RAxML tree from an alignment of the rDNA sequences (SSU+5.8s+LSU) of the newly assembled genomes. The same topology was obtained from an IQ-TREE analysis. Bootstrap values (1000 replicates) are shown above or next to the branches. The first bootstrap values correspond to the RAxML analysis and the second ones correspond to the IQ-TREE analysis. Full species names are presented in Table S1. Strain identifiers are included when more than one strain has the same species name in the complete set of species analyzed in this study (Table S1 and S2).

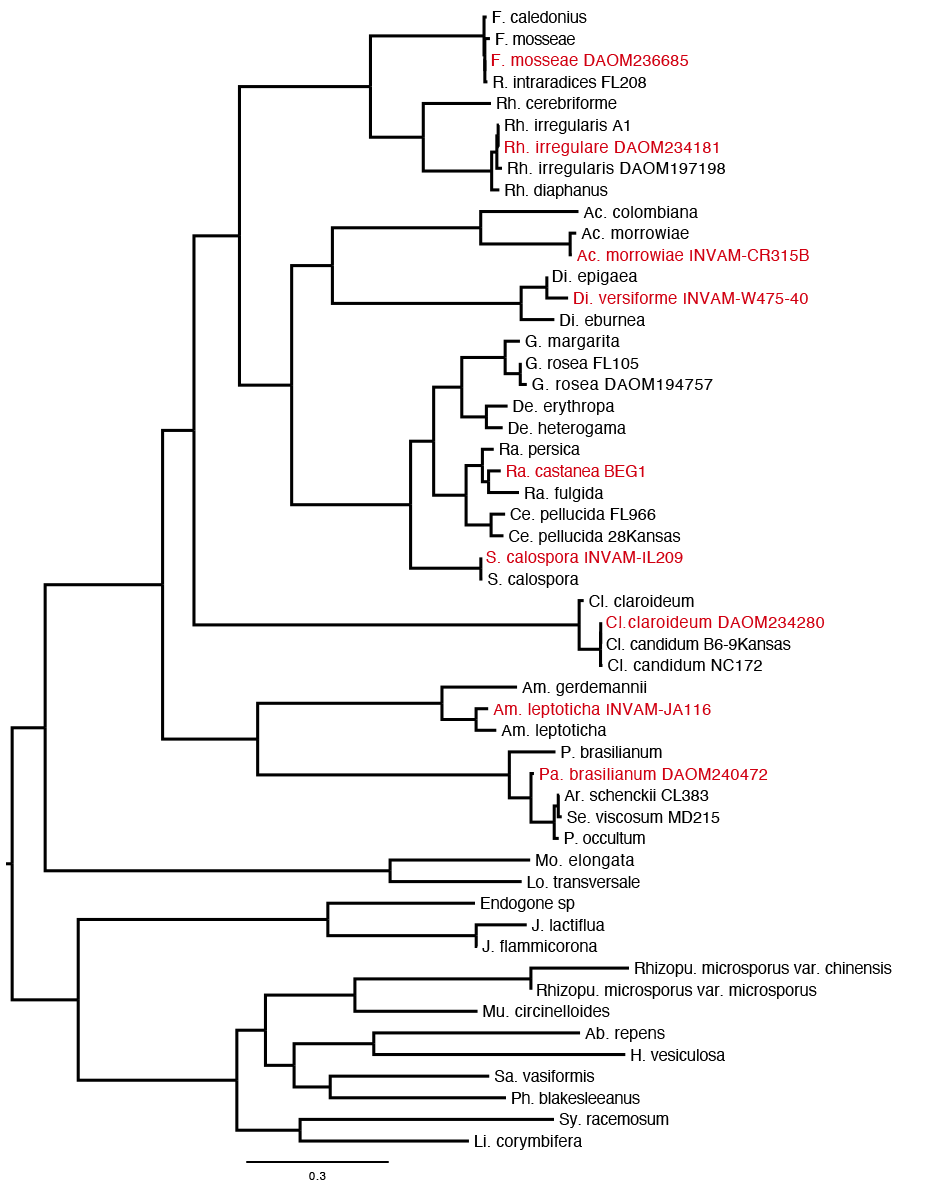

**Figure S3.** Best maximum likelihood RAxML tree from a concatenated alignment of 17 single copy orthologs shared among at least 50% of the taxa. Taxa in red show transcriptomic data from Beaudet *et al.*, 2018 and include species name and strain nr. The transcription data from Beaudet *et al.*, 2018 was not included in downstream analysis. Strain identifiers are included when more than one strain has the same species name in the complete set of species analyzed in this study (Table S1 and S2).

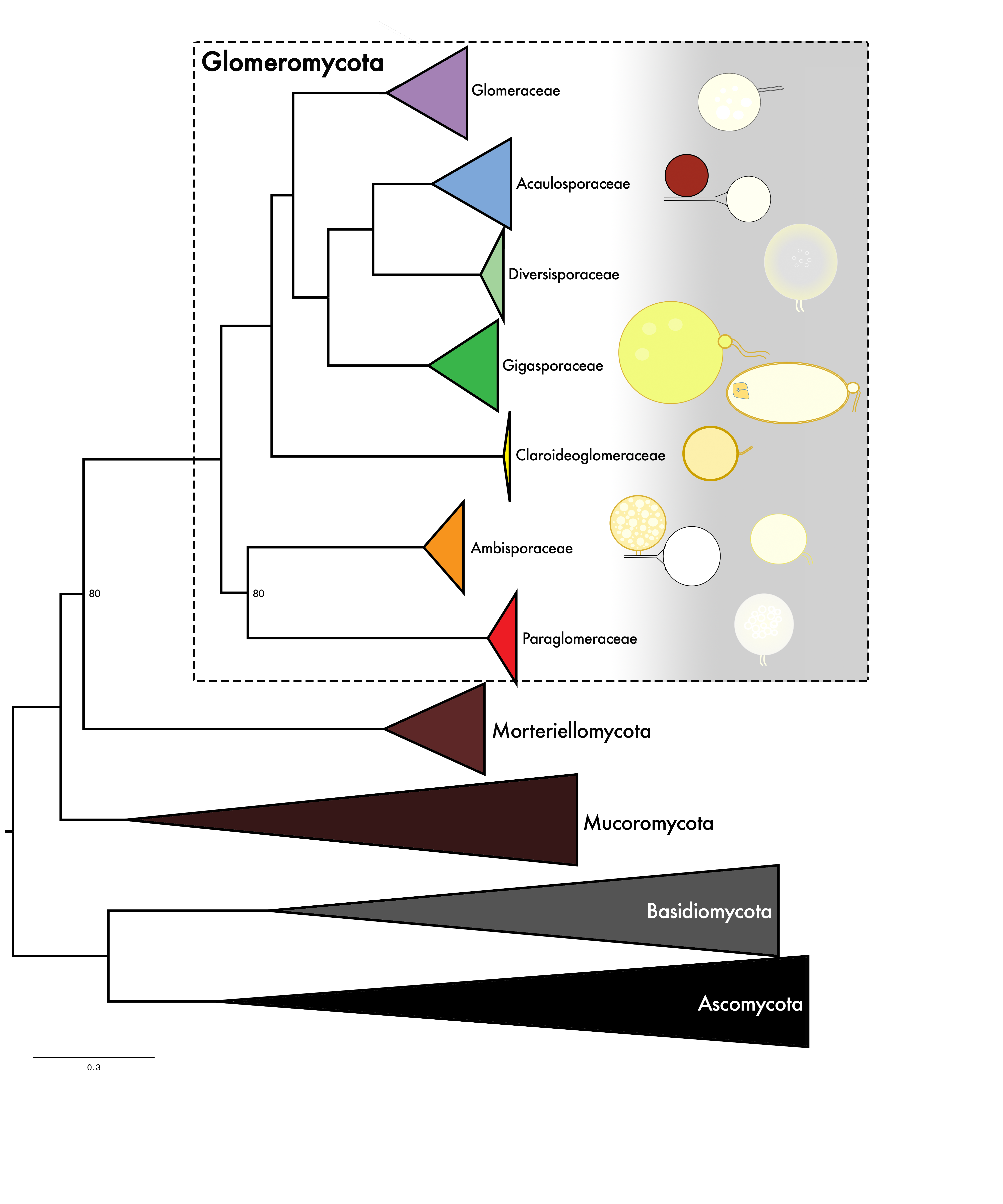

**Figure S4*.*** Best maximum likelihood RAxML tree from a concatenated alignment of 178 single copy orthologs shared among a minimum of 50% of the taxa. All branches have bootstrap support of 100 unless highlighted in the tree. With Dikarya represented by Basidiomycota and Ascomycota, as outgroup. Phyla are collapsed with the exception of Glomeromycota, where seven families are visualized in the dashed box. Typical spore morphologies are schematically illustrated to the right of each family and not drawn to scale. For expanded tree see Fig. S5a.

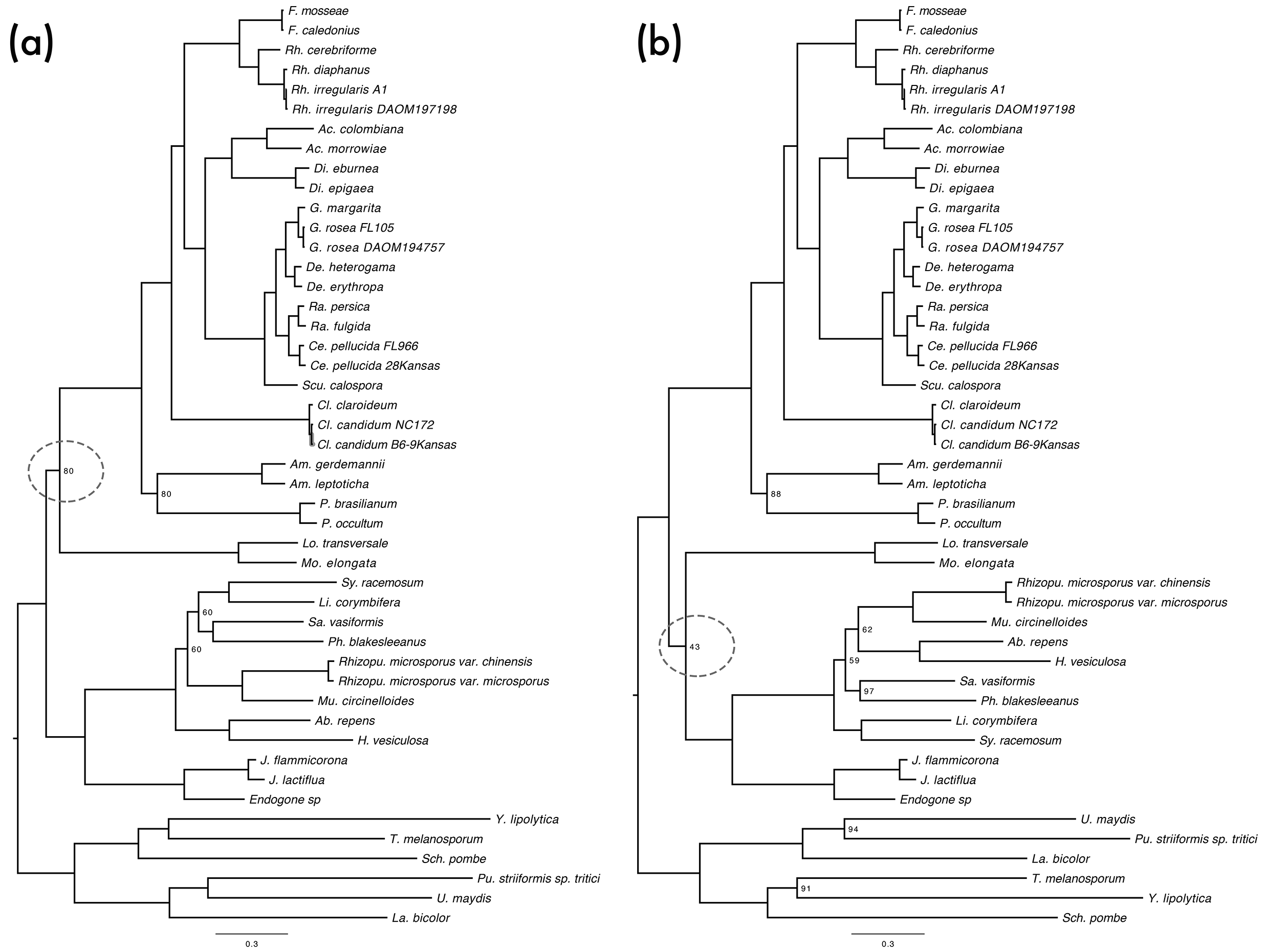

**Figure S5.** Maximum Likelihood Best Tree with RAxML (a), and IQ-TREE (b), from a concatenated alignment of 178 single copy orthologs shared among a minimum of 50% of the taxa. All branches have bootstrap support of 100 unless highlighted in the tree. With Dikarya as outgroup, represented by Basidiomycota and Ascomycota. Marked with a circle, we indicate the only conflict between these two topologies, in which Morteriellomycota appears as a sister group of Glomeromycota (a) or Mucoromycota (b). Full species names are presented in Table S1 and S2. Strain identifiers are included when more than one strain has the same species name in the complete set of species analyzed in this study (Table S1 and S2).

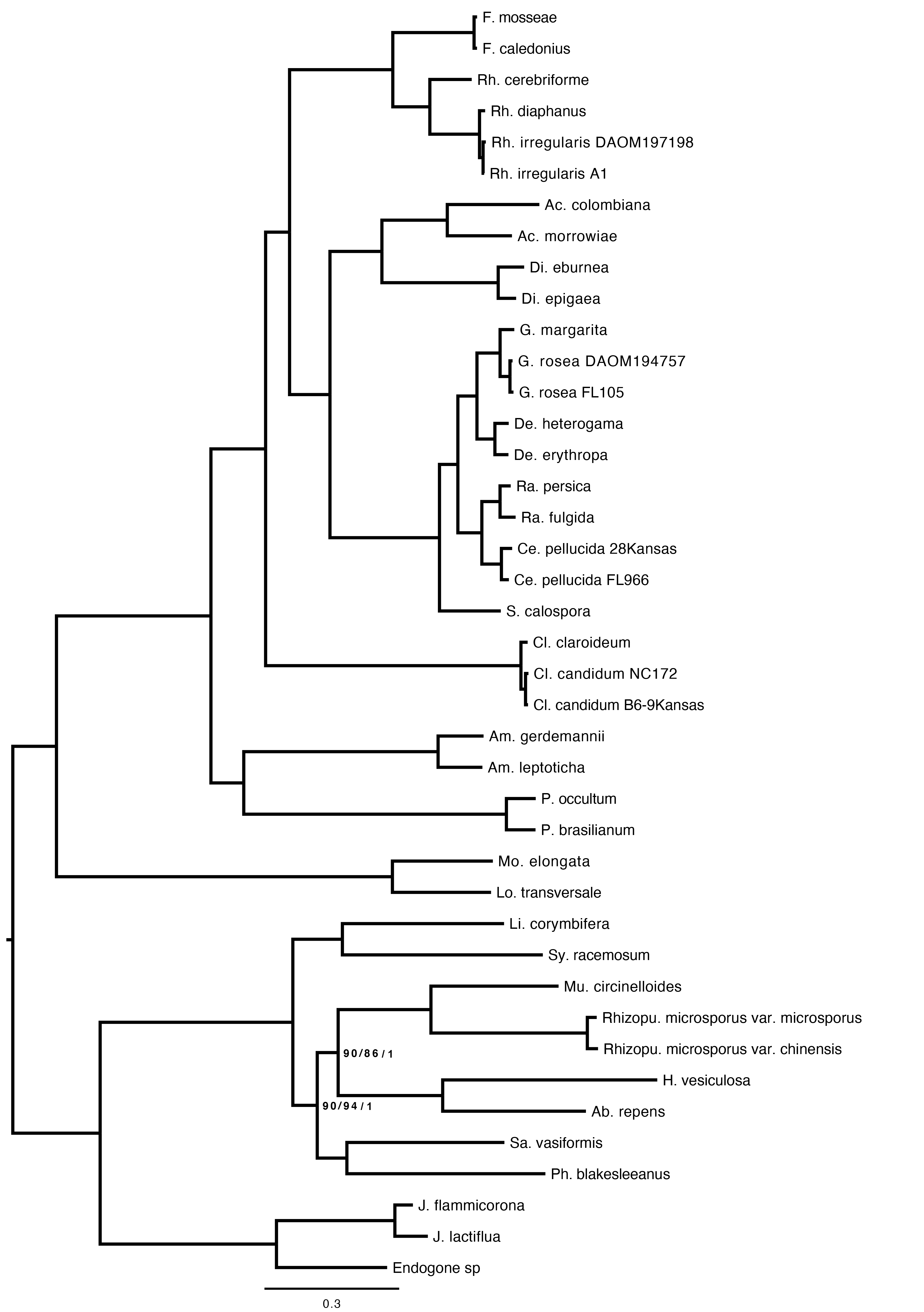

**Figure S6.** Expanded tree corresponding to Figure 1. Best ML tree inferred with RAxML from a concatenated alignment of 371 single copy orthologs shared among at least 50% of the taxa. Same topology was recovered using IQ-TREE and Bayesian inference. All branches have bootstrap support of 100 and posterior probabilities of 1 unless highlighted in the tree (RAxML/IQ-TREE/Bayesian). Strain identifiers are included when more than one strain has the same species name in the complete set of species analyzed in this study (Table S1 and S2).

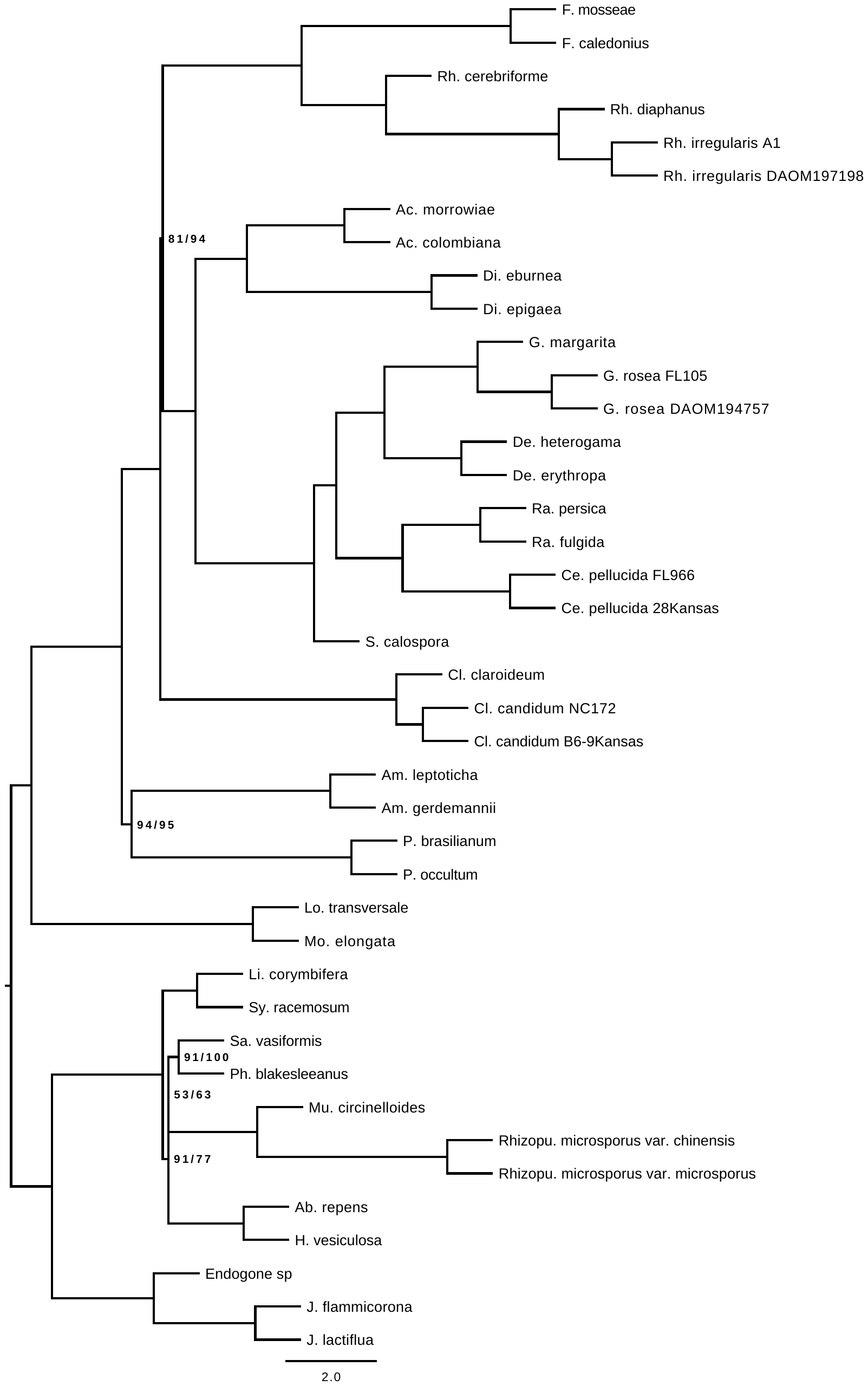

**Figure S7.** Multi-locus bootstrapping ASTRAL tree from 371 individual gene trees produced with RAxML and IQ-TREE. All branches have bootstrap support of 100 unless highlighted in the tree (RAxML/IQ-TREE). Strain identifiers are included when more than one strain has the same species name in the complete set of species analyzed in this study (Table S1 and S2).

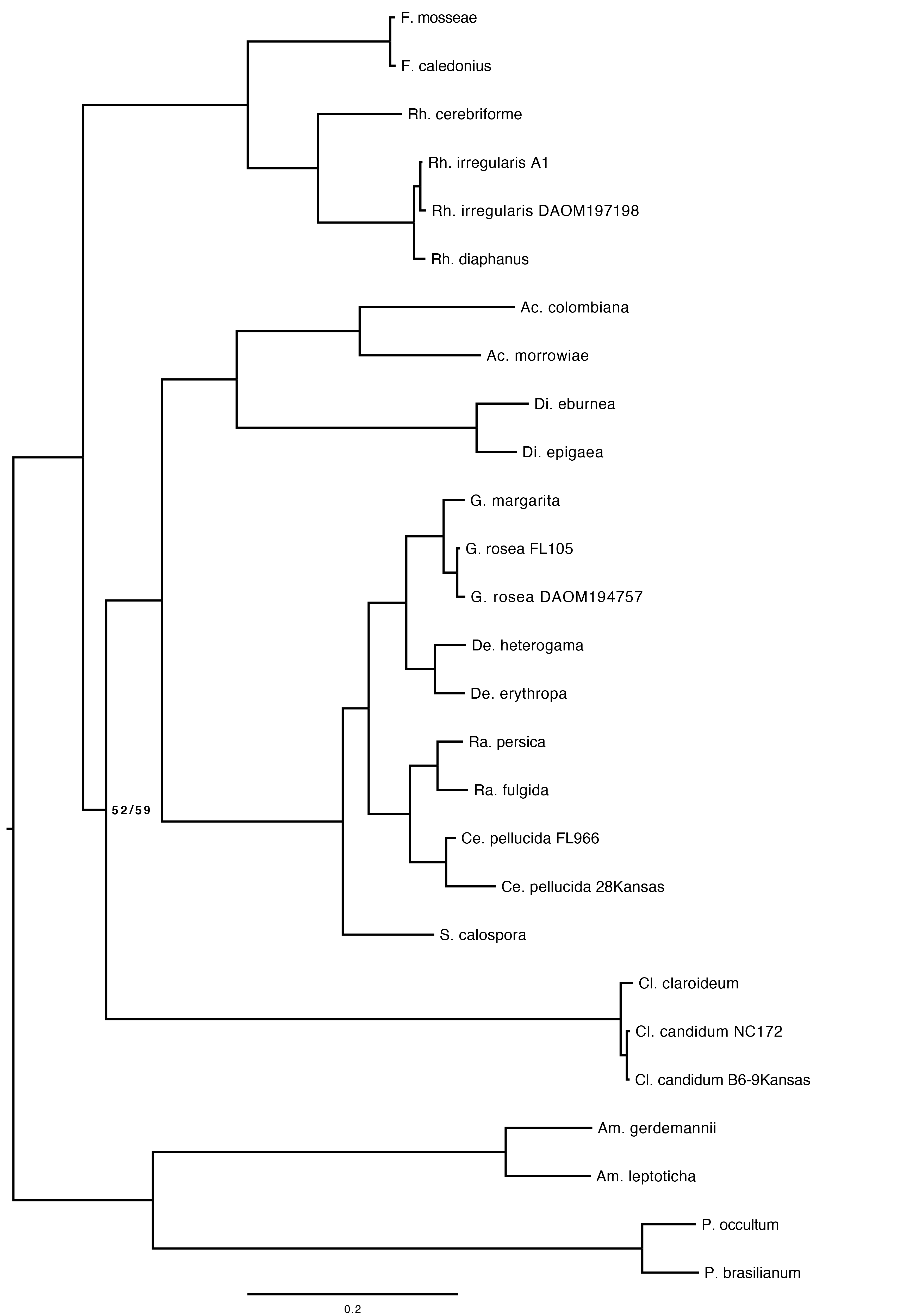

**Figure S8**. Best ML tree of Glomeromycota from 31 single copy orthologs shared among all taxa. Phylogenies produced with RAxML and IQ-TREE. All branches have bootstrap support of 100 unless highlighted in the tree (RAxML/IQ-TREE). Strain identifiers are included when more than one strain has the same species name in the complete set of species analyzed in this study (Table S1 and S2).

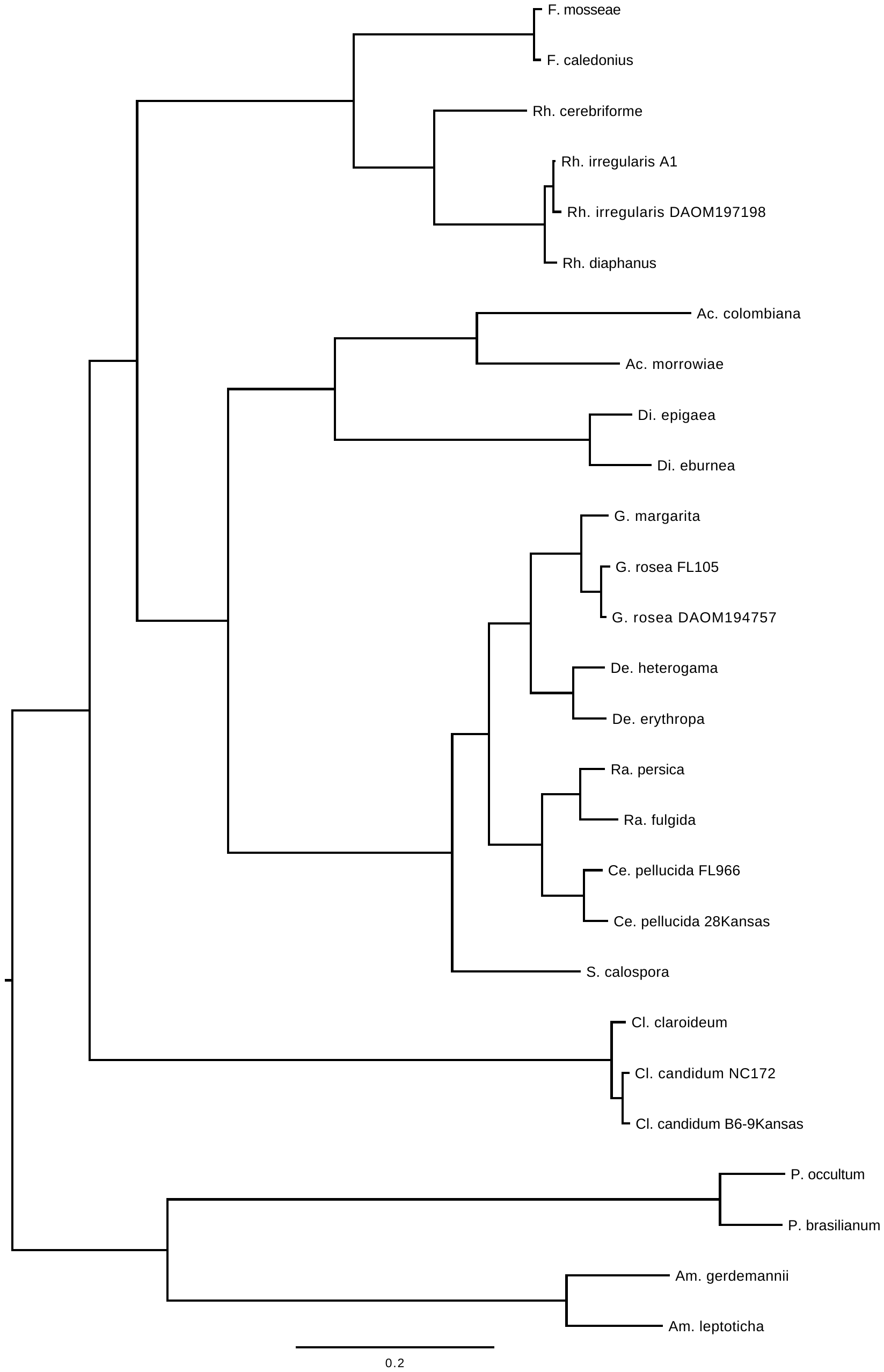

**Figure S9.** Best ML tree of Glomeromycota inferred from 1,737 single copy orthologs shared among, at least, 50% of the taxa. Phylogenies inferred with RAxML and IQ-TREE. All branches have bootstrap support of 100 in both analyses. Strain identifiers are included when more than one strain has the same species name in the complete set of species analyzed in this study (Table S1 and S2).

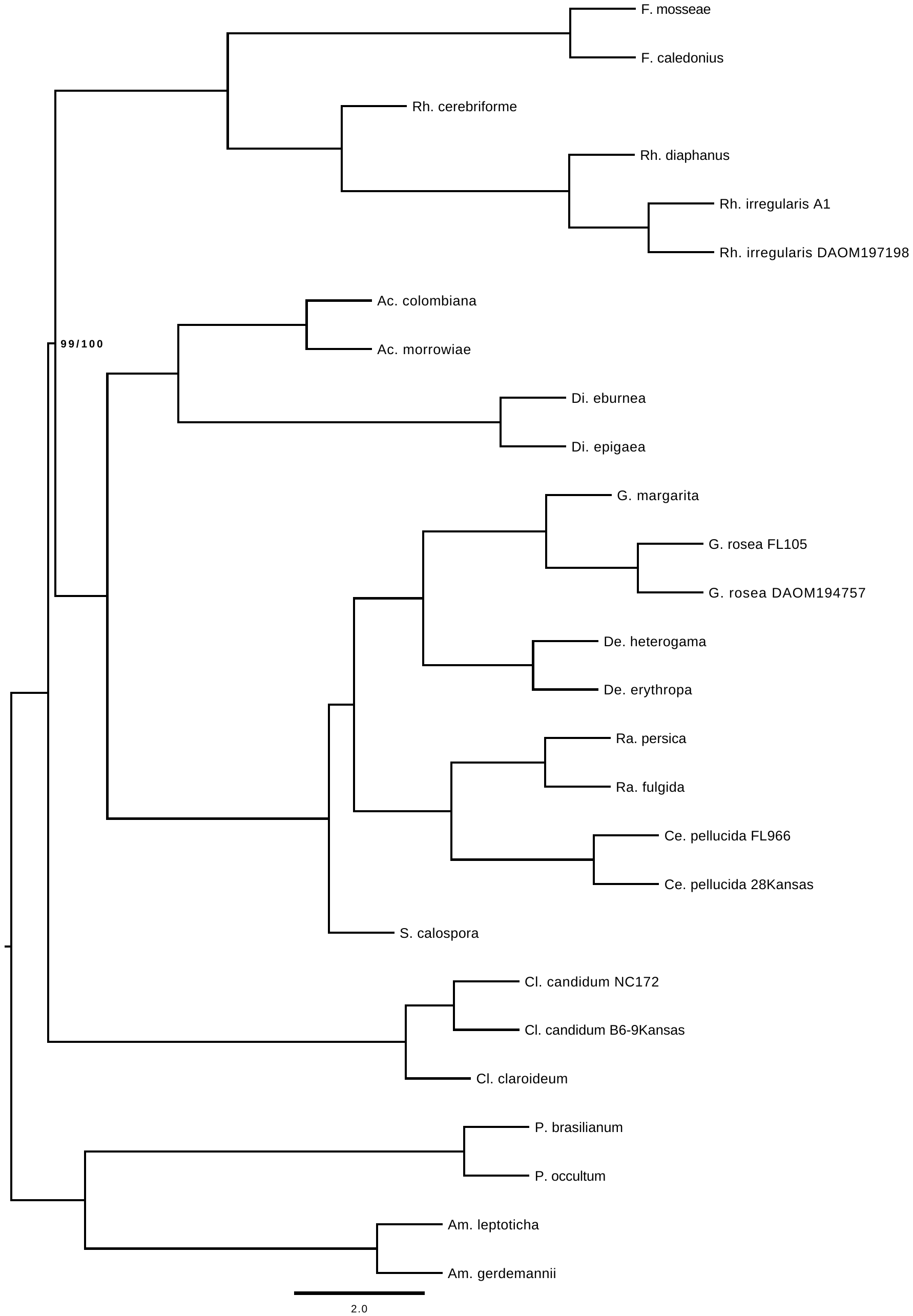

**Figure S10.** Multi-locus bootstrapping ASTRAL tree inferred from 1,737 individual gene trees produced with RAxML and IQ-TREE. All branches have bootstrap support of 100 unless highlighted in the tree (RAxML/IQ-TREE). Strain identifiers are included when more than one strain has the same species name in the complete set of species analyzed in this study (Table S1 and S2).

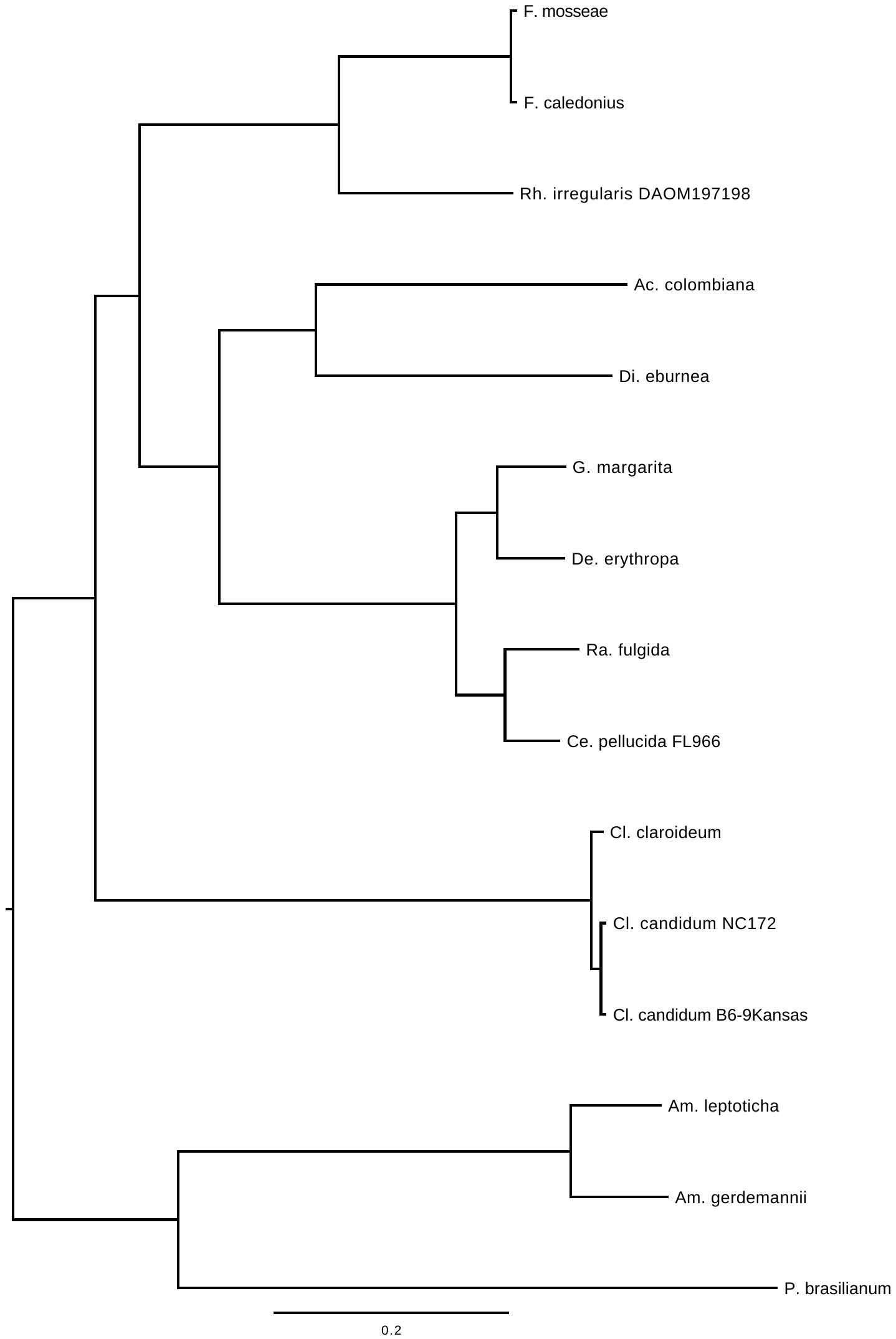

**Figure S11.** Best ML tree of Glomeromycota based on 799 single copy orthologs shared among 15 selected taxa (Table S4). Phylogenies produced with RAxML and IQ-TREE. All branches have bootstrap support of 100. Strain identifiers are included when more than one strain has the same species name in the complete set of species analyzed in this study (Table S1 and S2).

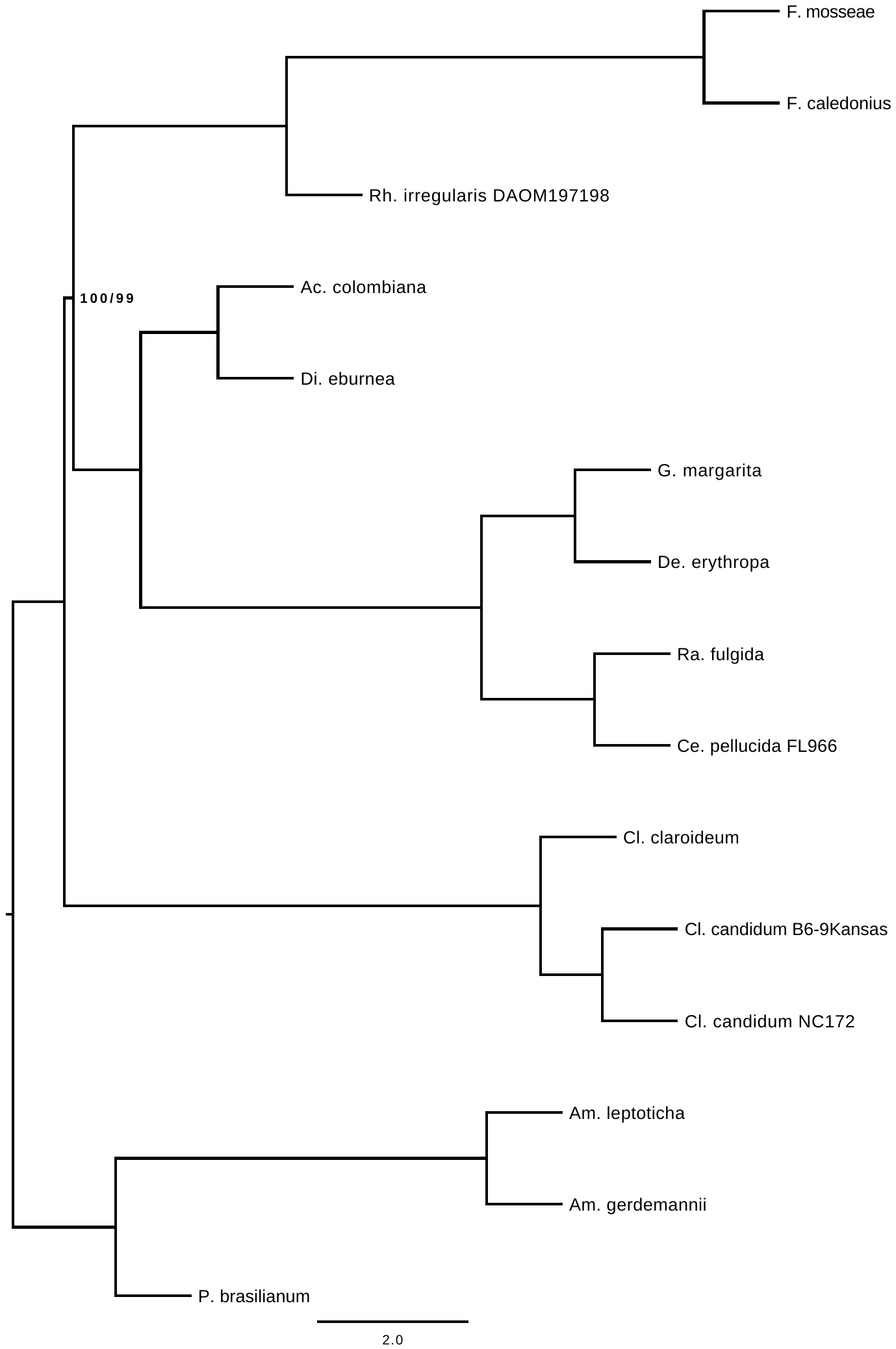

**Figure S12.** Multi-locus bootstrapping ASTRAL tree of 15 selected taxa of Glomeromycota (Table S4), inferred from 799 individual gene trees produced with RAxML and IQ-TREE. All branches have bootstrap support of 100 unless highlighted in the tree (RAxML/IQ-TREE). Strain identifiers are included when more than one strain has the same species name in the complete set of species analyzed in this study (Table S1 and S2).

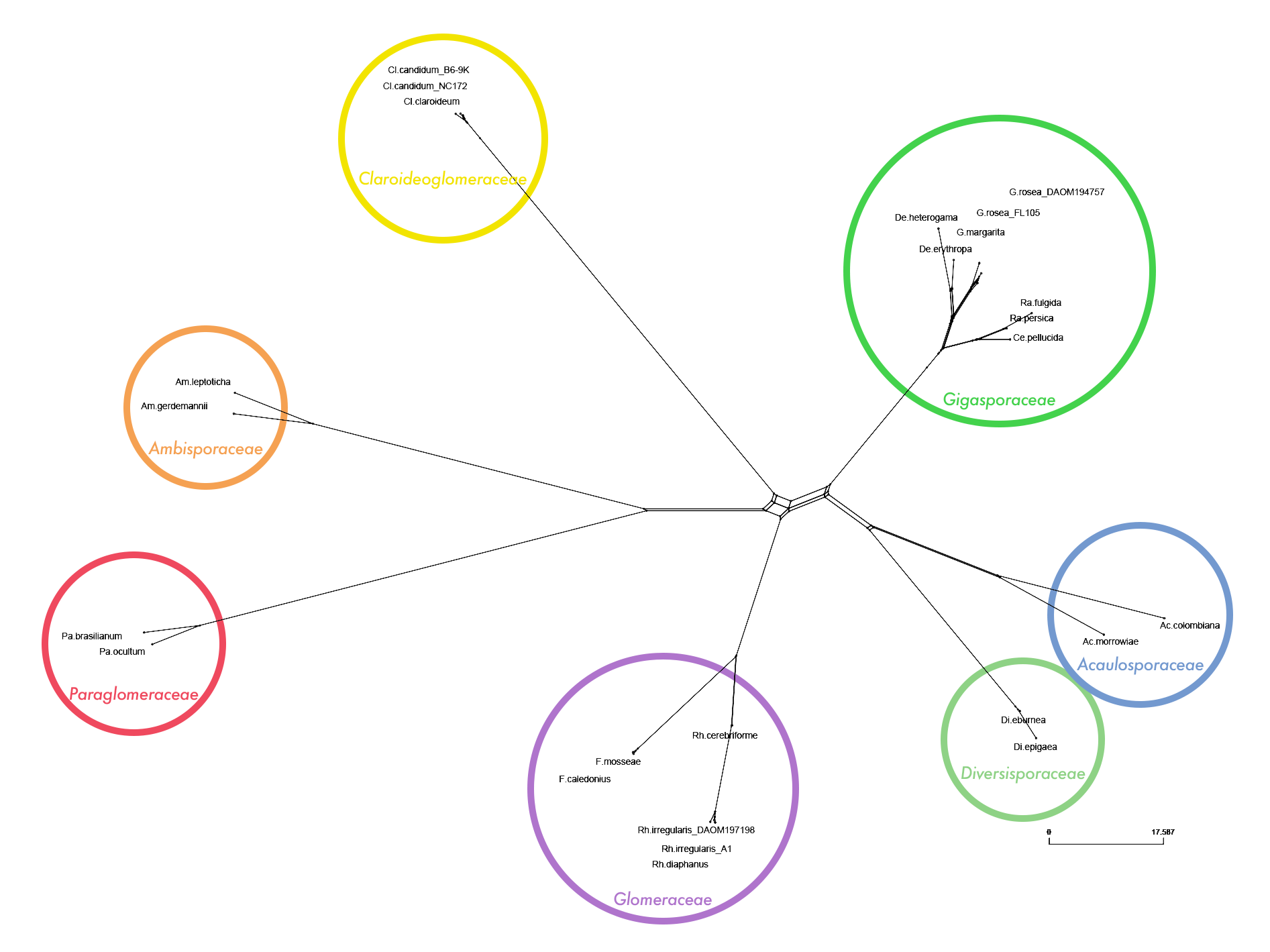

**Figure S13.** Expanded network from Figure 2a, in order to visualize the branch lengths between families (Circled and color coded according to Fig. 1 and Table S1). Produced with IQ-TREE network analysis and visualized in SplitsTree5 with maximum dimension splits filter of 2, using the dataset containing all Glomeromycota taxa, and 1,737 SCOs shared among, at least 50% of the taxa. Strain identifiers are included when more than one strain has the same species name in the complete set of species analyzed in this study (Table S1 and S2).

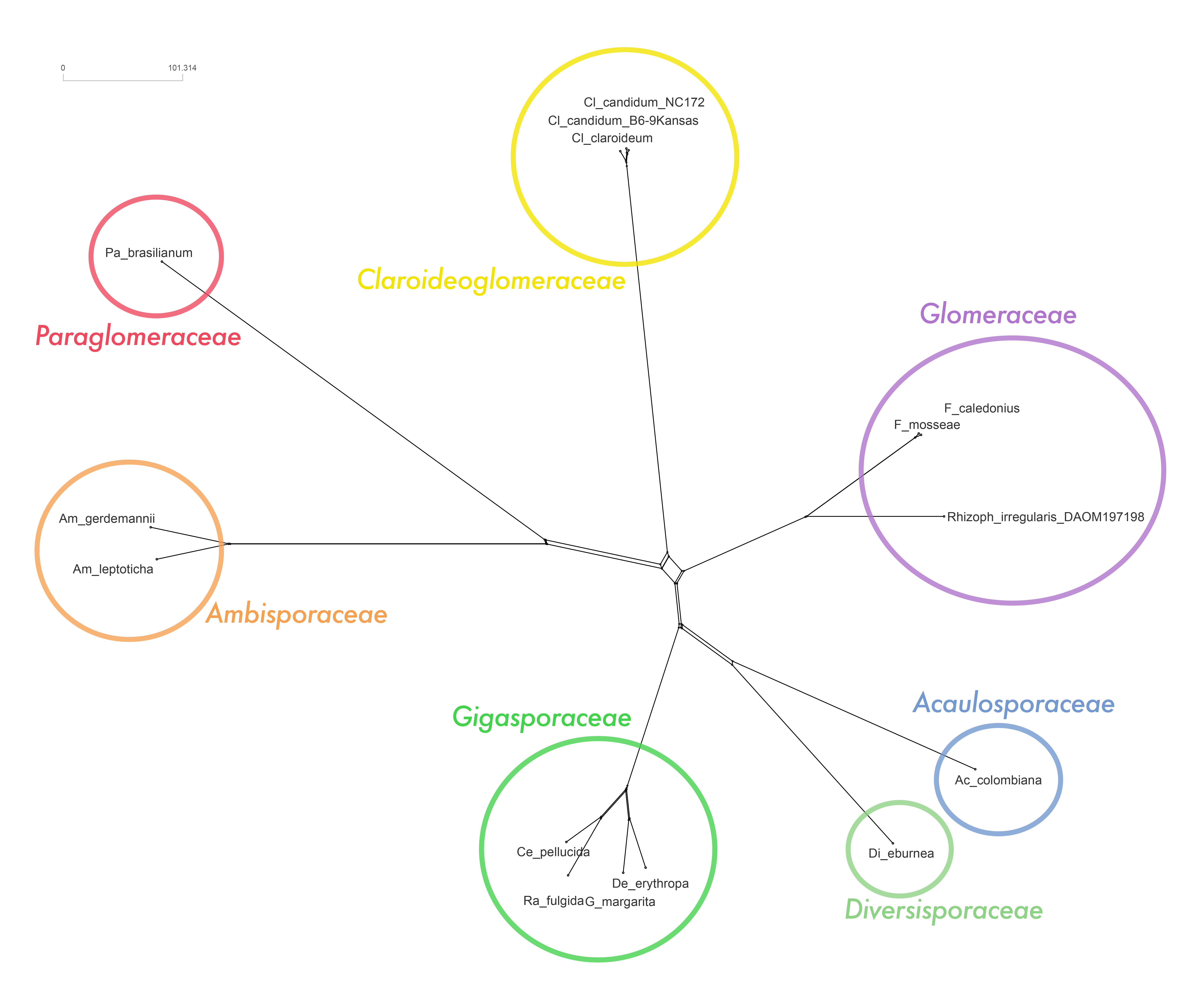

**Figure S14.** Network from IQ-TREE network analysis, using 799 single gene trees shared among 15 selected taxa (Table S4). Visualized in SplitsTree5, with a maximum dimension filter of 2. Full species names are presented in Table S1 and S2. Strain identifiers are included when more than one strain has the same species name in the complete set of species analyzed in this study (Table S1 and S2).

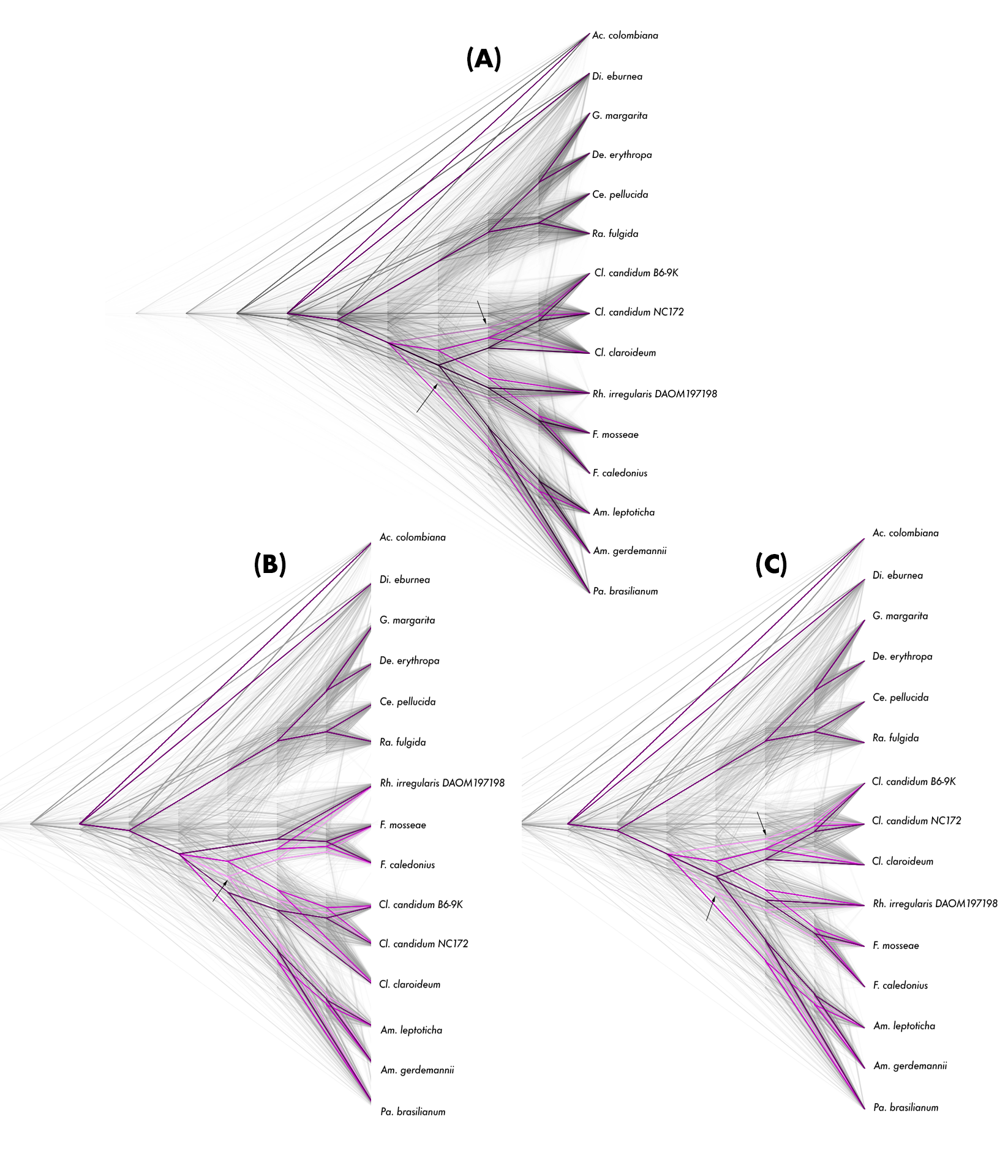

**Figure S15**. (A) Same Densitree as in Figure 2b based on 15 selected taxa, with order of taxa rearranged to visualize topology 3 (lightest shade of pink tree), in which Claroideoglomeraceae is sister group to Diversisporales (black arrows). (B) and (C) Densitree created with all the single gene trees produced with IQ-TREE. Full species names are presented in Table S1 and S2. Strain identifiers are included when more than one strain has the same species name in the complete set of species analyzed in this study (Table S1 and S2).

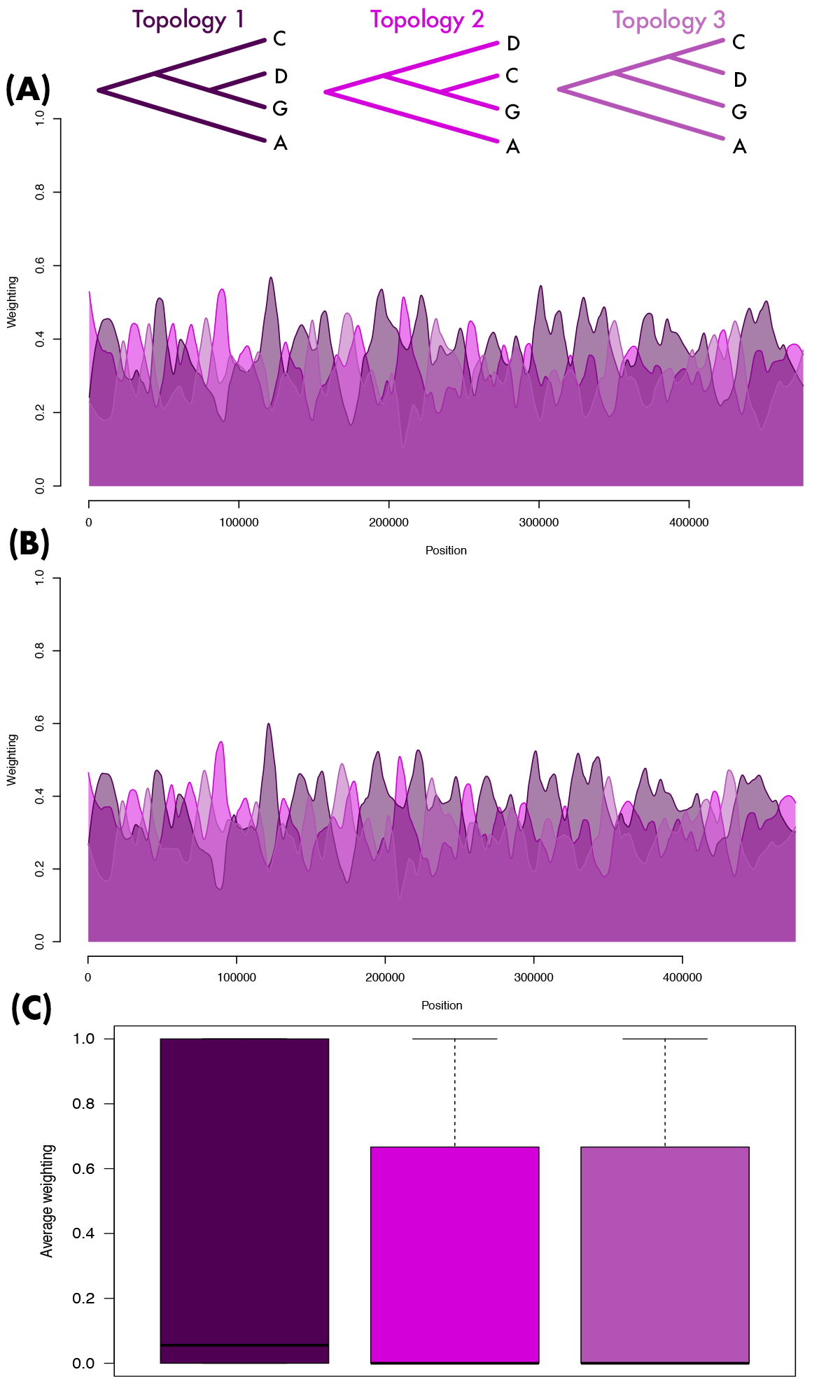

Figure S16. TWISST analysis on a selection of 15 taxa in Glomeromycota, grouped in four lineages: the families Claroideoglomeraceae (C), Diversisporales (D), Glomeraceae (G) and an outgroup with the two families Ambisporaceae and Paraglomeraceae (A), acting as taxa for topology analysis. Topology average weighting of 799 single gene tree topologies from best trees and 100 bootstrap trees for each best tree produced with RaxML (A), and IQTREE (B and C). On panels A and B, the concatenated alignment of all translated single copy orthologs corresponds to a total of almost 500k amino acids and is used as an artificial chromosome alignment. Data has been smoothed to windows of 50 aminoacids. Panel C shows the result of the average total weighting across all gene trees, produced with IQ-TREE (complementing figure 2c, produced with RaxML gene trees). Across all panels color coding indicates the weighted support for each of the three main topologies topology 1 (Darker shade of purple), topology 2 (bright pink) and topology 3 (lighter shade of purple).
